# A dynamic balance between neuronal death and clearance after acute brain injury

**DOI:** 10.1101/2023.02.14.528332

**Authors:** Trevor Balena, Kyle Lillis, Negah Rahmati, Fatemeh Bahari, Volodymyr Dzhala, Eugene Berdichevsky, Kevin Staley

**Author notes:** Correspondence: Kevin Staley, 114 16^th^ Street, Room 2600, Charlestown, MA, 02129.

## Abstract

After acute brain injury, neuronal apoptosis may overwhelm the capacity for microglial phagocytosis, creating a queue of dying neurons awaiting clearance. The size of this queue should be equally sensitive to changes in neuronal death and the rate of phagocytosis. Using rodent organotypic hippocampal slice cultures as a model of acute perinatal brain injury, serial imaging demonstrated that the capacity for microglial phagocytosis of dying neurons was overwhelmed for two weeks. Altering phagocytosis rates, e.g. by changing the number of microglia, dramatically changed the number of visibly dying neurons. Similar effects were generated when the visibility of dying neurons was altered by changing the membrane permeability for vital stains. Canonically neuroprotective interventions such as seizure blockade and neurotoxic maneuvers such as perinatal ethanol exposure were mediated by effects on microglial activity and the membrane permeability of apoptotic neurons, and had either no or opposing effects on healthy surviving neurons.

**Significance:** After acute brain injury, microglial phagocytosis is overwhelmed by the number of dying cells. Under these conditions, the assumptions on which assays for neuroprotective and neurotoxic effects are based are no longer valid. Thus longitudinal assays of healthy cells, such as assessment of the fluorescence emission of transgenically-expressed proteins, provide more accurate estimates of cell death than do single-time-point anatomical or biochemical assays. More accurate estimates of death rates will increase the translatability of preclinical studies of neuroprotection and neurotoxicity.

## Introduction

Neuronal cell death occurs on a temporal continuum with immediate, necrotic, or oncotic death at one end, and programmed or apoptotic death at the other (1–3). Apoptosis occurs both physiologically (4) and following traumatic and ischemic injuries (2, 5–7). When many neurons are injured simultaneously, the process of apoptosis is prolonged (8, 9), a condition referred to as delayed neuronal death (10). Apoptosis is irreversibly initiated by activation of proteolytic caspases that initiate the disassembly of the synthetic machinery of the cell (11, 12). Key features of apoptosis include early impairment of protein synthesis (13–15) but sustained ATP production despite caspase activation, DNA fragmentation, and increases in mitochondrial calcium levels (7, 16, 17). Signals leaking from the increasingly permeant membrane of apoptotic neurons attract microglia (18). Apoptosis ends in microglial engulfment and phagocytosis of the shrunken, pyknotic neuron (3, 19), a process termed efferocytosis (20, 21) that is used henceforth.

Immediate neuronal death is associated with mitochondrial failure, swelling, and cytoplasmic membrane rupture (22). Immediate cell death is difficult to assay microscopically because it is rapid and ends with cellular dissolution (2, 23). It is most often assessed non-cytologically by release of cytoplasmic enzymes such as lactate dehydrogenase (LDH) (24) or estimated from local changes in water diffusivity associated with cellular swelling (25, 26). Because of its rapid onset, immediate neuronal death has often already occurred at the time of clinical presentation, and is not studied extensively with respect to neuroprotection. In contrast, because apoptosis after injury occurs over much longer time spans (9, 27, 28), many cytological biomarkers have been developed to facilitate microscopic analysis of the number of dying cells (29). Although caspases are key initiators of apoptosis, their activity is transient (11), so assays of caspase activity are not widely used to quantify delayed neuronal death. Plasma membrane deterioration including blebbing and phosphatidylserine exposure occurs during apoptosis (30, 31). Accordingly the membrane permeability of normally-excluded dyes such as silver, fluorojade, ethidium bromide and propidium iodide (PI) are frequently used to identify neurons undergoing apoptosis (32–35).

The brain slice preparation is useful for the study of neuronal death after acute injury. Following the hypoxic-ischemic and traumatic injuries involved in brain slicing, both immediate (36) and delayed neuronal death (37, 38) are prominent. Surviving neurons in rodent organotypic hippocampal slice cultures retain their structural and electrophysiological integrity (39) and can be stably maintained in culture for many weeks (40), greatly facilitating the longitudinal assessment of neuronal death, for example by the ongoing emission of a fluorescent protein (FP) (41–43).

When the brain is injured, many neurons die in a short time frame. Efferocytosis of these neurons takes time (44, 45), so single-time-point analyses of neuronal death convolve several variables: the time since injury, the membrane permeability (i.e. staining properties) of neurons undergoing apoptosis, and the rates of microglial migration and efferocytosis. As an example of potential problems encountered when assessing neuronal death solely with vital dye assays, we present a study we published previously (38). Deafferented neurons in brain slice cultures develop extensive recurrent axonal sprouting (46) and consequently develop spontaneous electrographic seizures (47, 48) that peak in the second week in vitro; this coincides with the peak of neuronal staining with PI (38). In this preparation, seizures are associated with increased neuronal death (49), while agents that reduce neuronal activity and seizures sharply reduce peak PI staining (37, 38, 50). We interpreted these findings as evidence that recurrent seizures were inducing large numbers of neurons to undergo apoptosis and consequently stain with PI, with about 200 CA1 neurons per slice staining positive for PI on each imaging day. However, if 200 CA1 neurons die every day in a preparation with approximately 2000 CA1 neurons (51) there should be no neurons left in 10 days. Yet we found abundant healthy CA1 neurons in these cultures for over 6 weeks (38, 40). The abundance of surviving neurons suggests that it is not possible to estimate the rate of neuronal death solely with PI staining as PI may *overestimate* the rate of neuronal death, and also raises the possibility that seizure activity is affecting this *overestimate* rather than the survival of healthy neurons. For example, seizure-induced reductions in microglial efferocytosis of apoptotic neurons could increase the number of visibly dying neurons (52, 53). Alternatively, changes in the membrane permeability of dying neurons could alter the fraction of these neurons that stain for cell death biomarkers such as PI.

We therefore tested the degree to which changes in either microglial efferocytosis rates or the permeability of the membranes of neurons undergoing apoptosis altered the number of neurons that were positive for biomarkers of apoptosis following acute brain injury (Figure 1). These numbers were compared to the number of neurons that were known to be healthy based on the longitudinal assessment of the fluorescence of transgenically expressed cytoplasmic proteins using serial two-photon neuronal imaging (41–43).

**Fig. 1.**
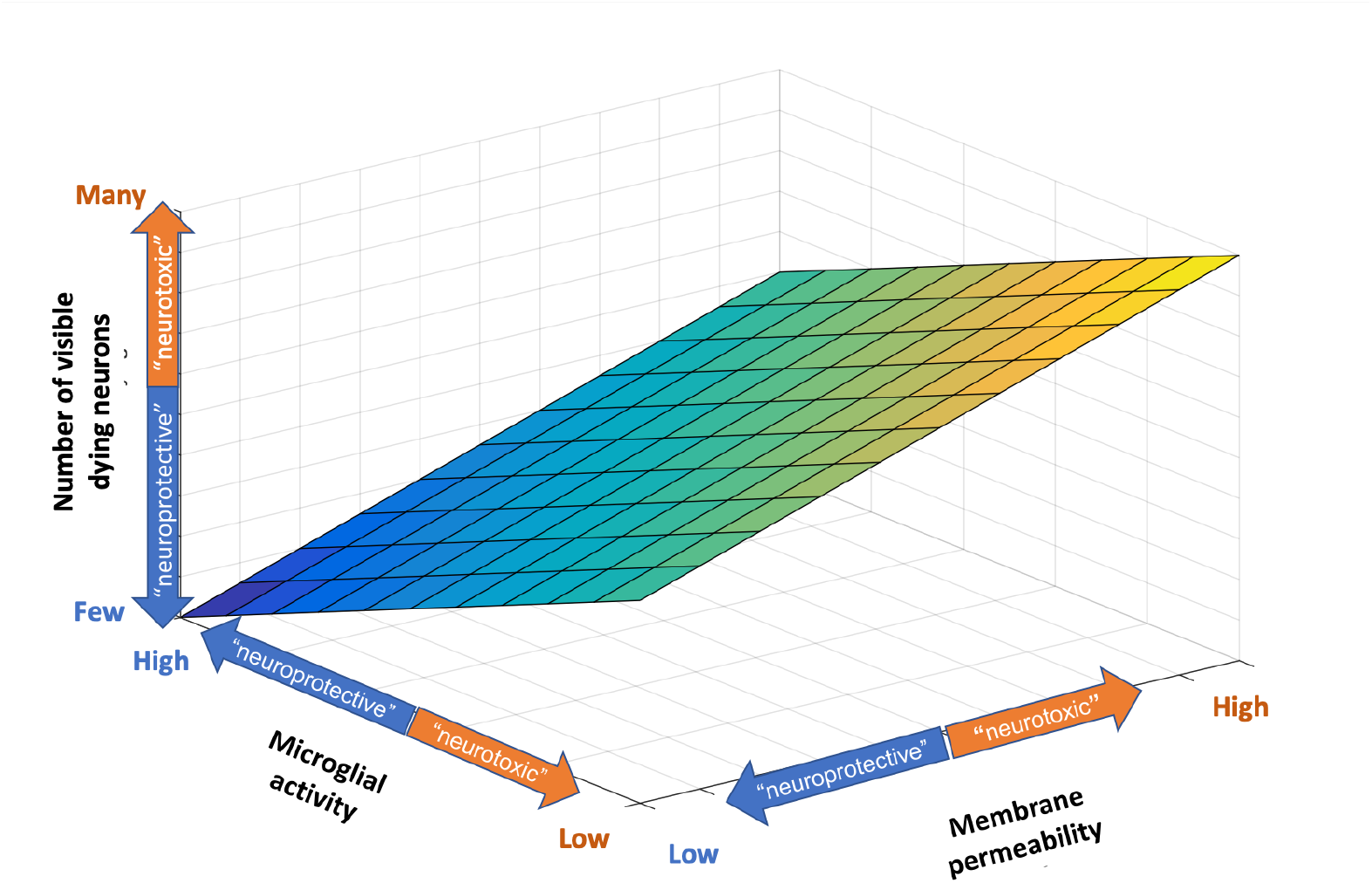
Apparent neuroprotective and neurotoxic effects due to changes in membrane permeability or the rate of microglial engulfment of apoptotic neurons. If the principal assay is the number of visibly dying neurons, which is plotted on the z-axis, then interventions that alter this number can be interpreted as neuroprotective or neurotoxic, independently of the effects of these interventions on neuronal death. For example, interventions that alter the rate of microglial efferocytosis of dying neurons may increase or decrease the number of visibly dying neurons, and could therefore be interpreted as neurotoxic or neuroprotective. Similarly, interventions that alter neuronal membrane permeability may change the degree of neuronal staining with biomarkers of cell death, which would also alter the number of visibly dying neurons. Such an effect would also be interpreted as neurotoxic or neuroprotective independently of an effect on the rate of neuronal death.

## Results

### Cell death stains and fluorescent protein expression

We considered 2 possible explanations for the apparent over-reporting of neuronal apoptosis by PI staining in our prior study (38). First, PI might also be staining healthy neurons. Healthy neurons can be assayed by robust emission of transgenically expressed fluorescent proteins (41–43, 54, 55). To address potential differences in labeling by the PI vs fluorescent protein biomarkers, slices were used from wild type (WT) mice in which neurons expressed green fluorescent protein (GFP) following infection of slices with the adeno-associated virus (AAV) AAV9-hSyn-eGFP vector. Slices were also used from Clomeleon mice in which neurons transgenically expressed both cyan fluorescent protein (CFP) and yellow fluorescent protein (YFP) under control of the Thy1 promoter (56); henceforth Clm will be used to refer collectively to the fluorescent proteins expressed in slices from Clomeleon mice. These slices were stained with PI on days *in vitro* (DIV) 5-7 to assess the degree of overlap (Figure 2A). Neurons with bright emission from transgenic fluorophores had normal morphology (Figure 2B) and minimal nuclear PI staining (Figure 2C).

**Fig. 2.**
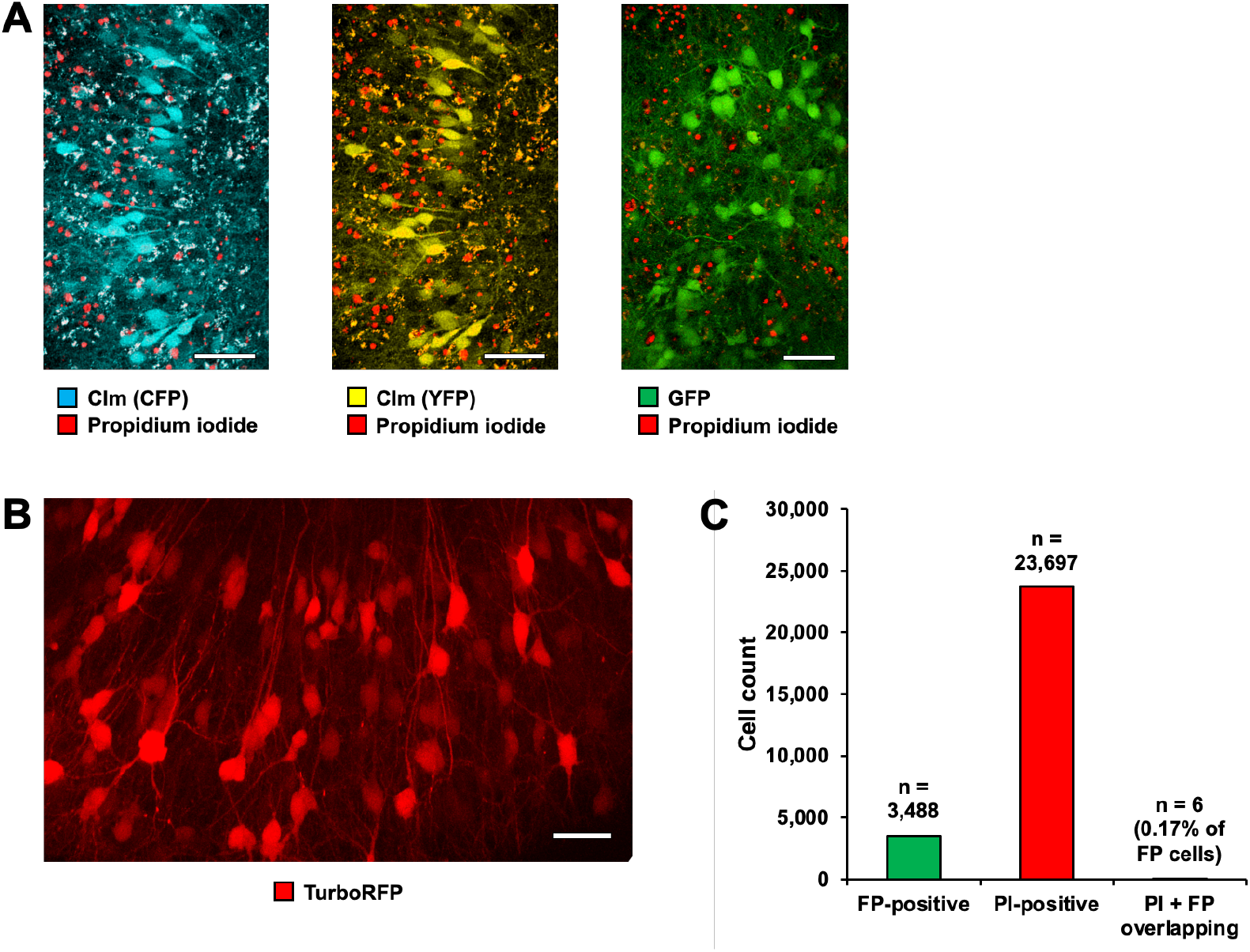
Overlap of transgenic fluorescent protein emission and propidium iodide nuclear staining. **(A)** Visualization of the lack of overlap between propidium iodide-positivity and FP-positivity. In slices made from Clomeleon mice imaged at DIV 7, neither the CFP emission (left) nor the YFP emission (middle) of Clm was present in cells with PI-positive nuclei. The same was true of cells in WT slices that expressed GFP (right), imaged at DIV 5. Scale bars = 50 μm. **(B)** Morphology of CA1 neurons in organotypic hippocampal slice cultures imaged at DIV 11. Neurons with bright emission from expressed TurboRFP (FP+; see Methods) demonstrate normal somata, nuclei, and dendritic arbors. Scale bar = 50 μm. **(C)** FP+ neurons are propidium-iodide negative. Across 23,697 PI-positive neurons and 3,488 GFP-positive neurons, only 6 of the GFP-positive neurons (or 0.17%) also exhibited PI-positivity. n = 12 slices.

Between 23,697 PI-positive cells and 3,488 GFP-positive neurons identified by the ImageJ plugin TrackMate, only 6 instances of overlap (or 0.17% of GFP-positive neurons) were seen (n = 12 slices). In less healthy slices, overlapping fluorescence in the PI and fluorescent protein emission bands were more common due to the wide-spectrum emission properties of autofluorescent cellular debris; still, between 265,382 PI-positive cells and 53,750 GFP- or Clm- positive neurons identified by ImageJ, only 2,434 instances of overlap (or 4.53% of FP-positive neurons) were seen when these slices were included (n = 99 slices). Less healthy slices could be identified by the presence of autofluorescent debris, which was significantly smaller than healthy neurons. This reduced the average diameter of cells identified by CellProfiler (18.71 ± 0.26 μm vs. 16.98 ± 0.14 μm, p < 0.0001, n = 12 vs. 87 slices) as well as the area (341.50 ± 9.08 μm^2^ vs. 275.30 ± 4.48 μm^2^, p < 0.0001, n = 12 vs. 87 slices) and perimeter (87.62 ± 1.12 μm vs. 76.22 ± 0.68 μm, p < 0.0001, n = 12 vs. 87 slices) in less healthy slices compared to healthy slices.

The number of FP+ neurons was 15-20% of the number of PI+ neurons in these preparations, which is to be expected since PI stains the vast majority of late-stage apoptotic neurons but fluorescent protein expression (be it viral or transgenic) may only be detected in a fraction of healthy neurons (40). These results indicate that FP-positive neurons are healthy, with intact cellular and nuclear membranes that do not admit PI. They also confirm that a considerable number of neurons are undergoing apoptosis in these organotypic slice cultures days or weeks after the initial brain slicing injury.

There was also minimal overlap between neurons with robust emission from transgenic fluorescent proteins and neurons exhibiting elevated caspase activity. Slices in which neurons expressed the red fluorescent protein (RFP) TurboRFP were loaded with fluorochrome-labeled inhibitors of caspases (FLICA) to visualize caspase activity. Starving these slice cultures of nutrients and oxygen by stopping artificial cerebrospinal fluid (ACSF) perfusion resulted in quenching of neuronal TurboRFP fluorescence emission after 1-2 hours of starvation. FLICA signals were elevated in many neurons within the same time frame (Figure 3A). In some cases, TurboRFP quenching and increases in FLICA occurred in the same neurons with a period of overlap of less than 1 hour (the interval between sequential imaging sessions). Subsequently, FLICA fluorescence also decreased, indicating that the elevated caspase activity is transient (Figure 3B) (9, 11). Indeed, instances of overlap between FLICA and fluorescent protein emission were rare; in a 590×590×140 (X:Y:Z) μm region of a normally perfused slice, 107.7 ± 49.0 TurboRFP-positive neurons, 41.6 ± 16.8 FLICA-positive cells, and 12.9 ± 8.6 cells that were positive for both could be seen (Figure 3C), with only 6.5 ± 3.3% of TurboRFP-positive cells also being FLICA-positive, and only 17.0 ± 9.6% of FLICA-positive cells also being TurboRFP-positive (Figure 3D) (n = 754 TurboRFP-positive neurons and 291 FLICA-positive neurons in 7 fields of view from 3 slices). This same lack of overlap was also seen when TurboRFP- and FLICA-positive cells were imaged serially for several hours, with TurboRFP-positivity dropping and FLICA-positivity increasing shortly after apoptosis was induced by stopping perfusion. After 6 hours of starvation both fluorescent protein emission and FLICA fluorescence had diminished, though some regions required a longer period of starvation than others (Figure 3E). Most FLICA+ cells were neurons because the fraction of newly FLICA+ cells that were currently, or had been previously, TurboRFP+ was comparable to the typical fraction of neurons that label with AAV9-synapsin-driven fluorescent proteins in this preparation (Figure 3F) (40). These experiments indicate that neurons with bright emission from transgenic fluorescent proteins rarely stain for caspase activity or PI. We conclude that neurons expressing fluorescent proteins are healthy in this preparation, and that PI is not falsely labeling healthy neurons, so that the over-reporting of neuronal death by PI is not due to false positive PI staining of healthy neurons.

**Fig. 3.**
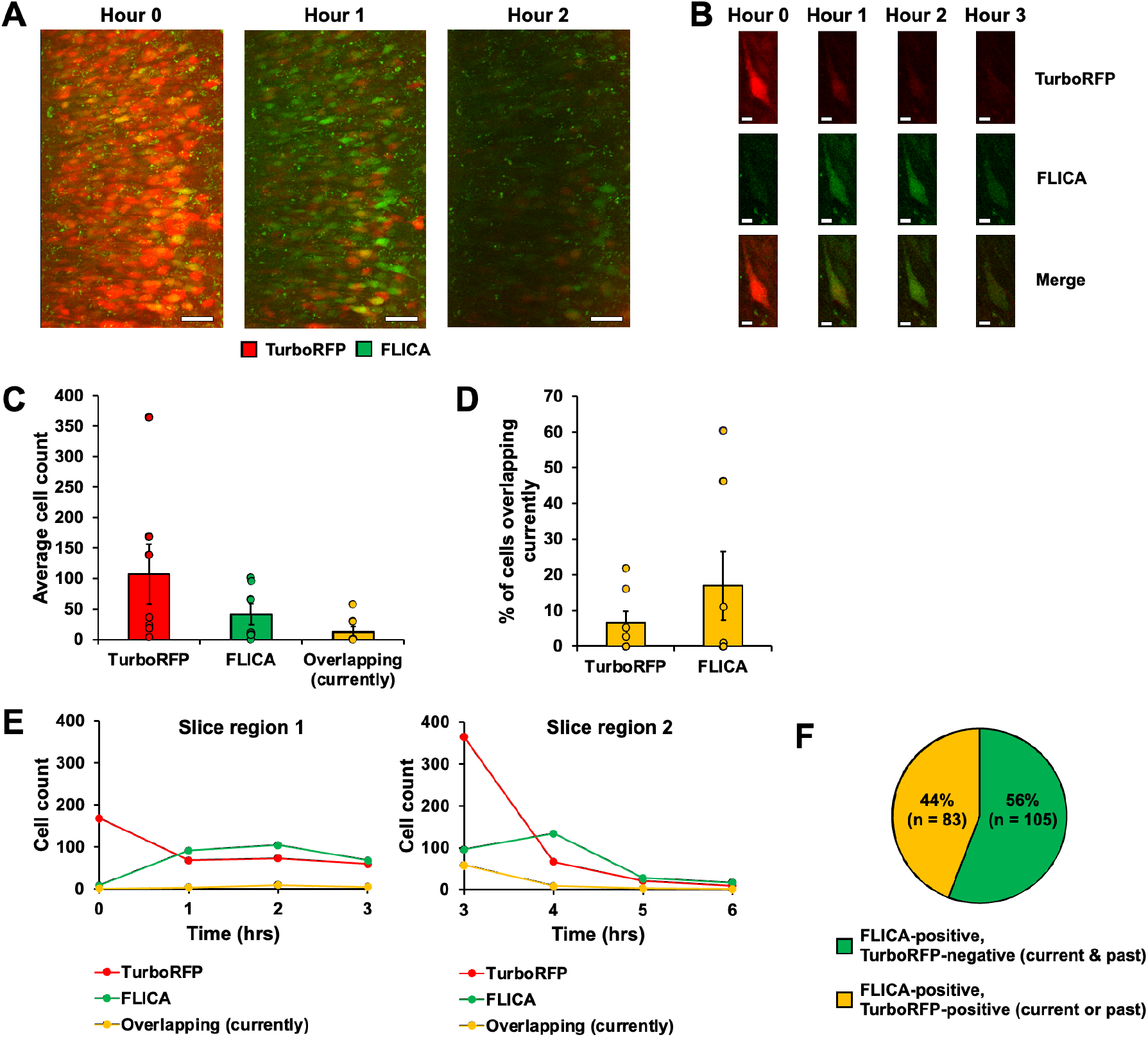
Overlap of transgenic fluorescent protein expression and FLICA. **(A)** Quenching of fluorescent proteins and an increase in caspase activity occur concurrently. Depriving CA1 neurons in an organotypic slice culture on DIV 7 of nutrients and oxygen by stopping perfusion at Hour 0 caused TurboRFP-positive neurons with little FLICA intensity (left) to become largely TurboRFP-negative and FLICA-positive (right) by Hour 1. By Hour 2, the TurboRFP-positivity has decreased even further and the transient caspase activity has largely ceased. Scale bars = 50 μm. **(B)** Example of transient overlap of fluorescent protein expression and increased caspase activity. On DIV 7 the neuron had strong TurboRFP expression and negligible FLICA signal at Hour 0 when perfusion was shut off. By Hour 1, both fluorophores could be seen in the same neuron; TurboRFP expression was substantially decreased but still visible, and FLICA emission was now strong. By Hour 2, TurboRFP expression was nearly gone and FLICA remained bright. But by Hour 3, the transient caspase activity had begun to decrease and the neuron was barely visible with either fluorophore. Scale bar = 10 μm. **(C)** FP + FLICA overlap is rare. Under control conditions with adequate perfusion, a 590×590×140 section of a slice contained 107.71±49.01 TurboRFP-positive neurons, 41.57±16.79 FLICA-positive cells, and 12.86±8.60 cells that were positive for both. n = 754 TurboRFP-positive neurons and 291 FLICA-positive neurons in 7 fields of view from 3 slices. **(D)** Few FP-positive cells exhibit elevated caspase activity. In the same slices as in (C), only 6.52 ± 3.32 % of TurboRFP-positive cells were also FLICA-positive. Conversely, 16.95 ± 9.63 % of FLICA-positive cells were also TurboRFP-positive. **(E)** Examples of FP-positivity and FLICA-positivity, and overlap over time. In two regions of a slice, imaged sequentially, TurboRFP-positivity dropped and FLICA- positivity increased once the perfusion was shut off but the amount of overlap at any given time remained low. After several hours the transient caspase activity decreased as well, lowering the FLICA-positivity. **(F)** Most FLICA-positive cells are neurons. Using TrackMate analysis of a 4-hour time series experiment, 44% of 188 FLICA-positive neurons were either currently also TurboRFP-positive, or had previously been TurboRFP-positive, which is very close to the rate of expression of TurboRFP in neurons in this preparation (40).

### Rates of apoptosis and efferocytosis

An alternate explanation for the over-reporting of apoptosis by PI staining is illustrated in Figure 4. The number of PI+ neurons in the preparation at any time point depends on the difference in rates of entry and exit into the pool of PI-receptive neurons, as well as the duration of that difference in rates. We therefore sought to characterize these rates. We first assayed the rate at which neurons *exited* the PI-receptive pool. Slices were prepared from Clomeleon mice (56) and on DIV 7, PI was added to the slices’ culture media. Following this one-time PI exposure, both PI and fluorescent proteins were visualized with two-photon microscopy on DIVs 7, 10, and 13, i.e. on the day of PI exposure and 3 and 6 days later. In the 12 slices the number of PI+ neurons declined monoexponentially with a time constant (𝛕) (𝛕 = 2.16 days, initial PI count = 967.0 ±106.1, n = 12 slices). These findings indicate that in hippocampal organotypic slice preparations on DIV 6-13, PI+ neurons waited an average of 1.5 days to be phagocytosed (Figure 5A). After 3 days, the percentage of PI-positive neurons remaining was negatively correlated with the number of visible FP+ neurons (R-squared = 0.5123, p = 0.0089, n = 12) (Figure 5B). The number of FP+ neurons should be larger in healthier slices (43). The monoexponential decline of PI+ neurons would be consistent with a first order process (57) that is limited by the number of PI+ neurons (Figure 5A). In other words, under these experimental conditions the rate of efferocytosis of PI+ neurons is proportional to the number of PI+ neurons in the apoptotic pathway.

**Fig. 4.**
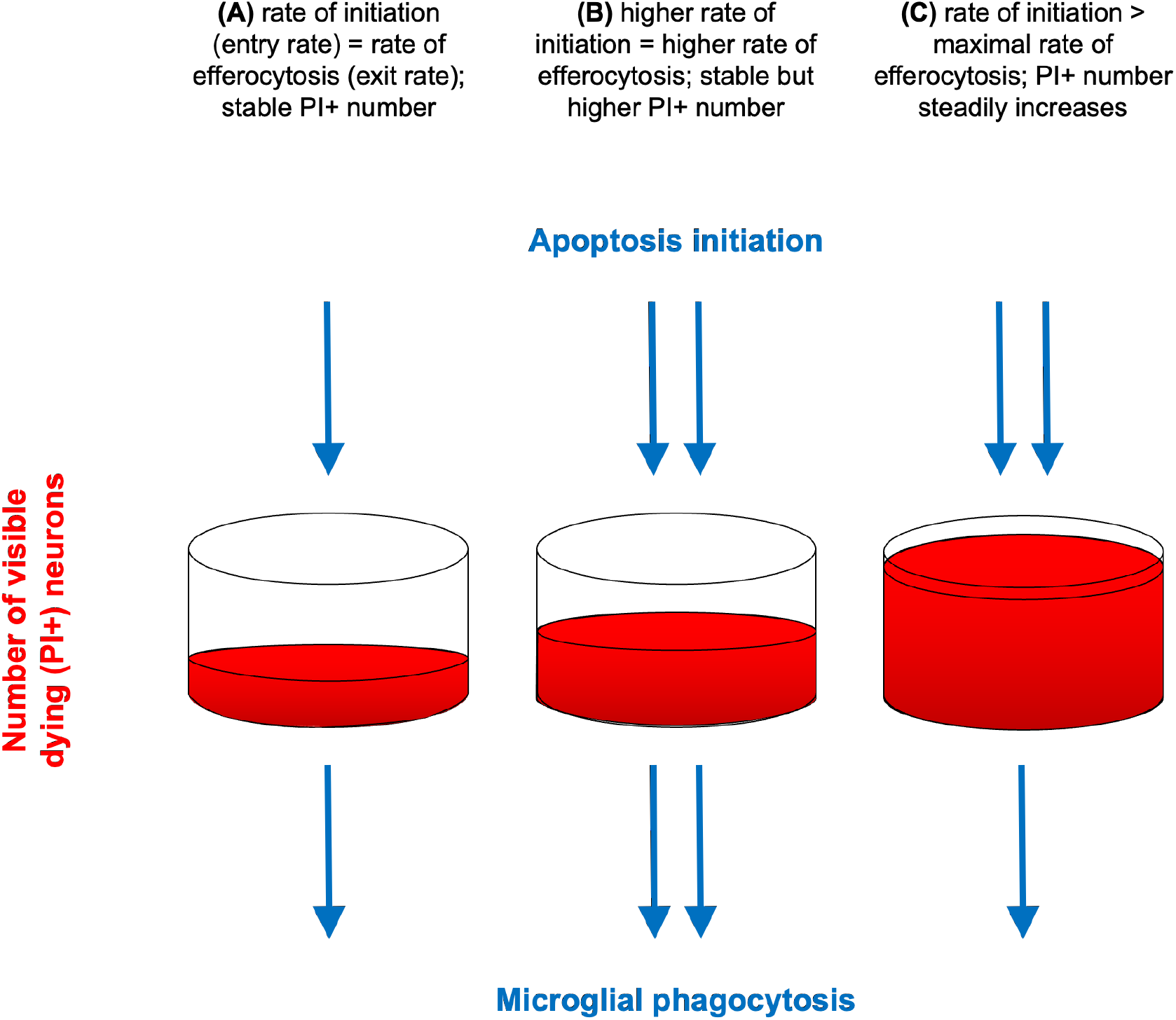
The size of the apoptosis biomarker-receptive pool depends on time, and the rates of apoptosis and efferocytosis. The fluid in the reservoir represents the number of visibly dying neurons, i.e those labeled by one-time staining with PI. **(A)** The number of visibly dying (PI+) neurons depends on both the rate of apoptosis and the rate at which PI+ neurons are phagocytosed. **(B)** If the rate of apoptosis doubles, the corresponding increase in the number of PI+ neurons also depends on how the rate of efferocytosis changes with the number of PI+ neurons. **(C)** If the rate of apoptosis increases beyond the maximum rate of efferocytosis, then the number of visibly dying neurons will steadily increase. Such an increase in apoptosis is sufficient but not necessary to increase the number of visibly dying neurons, because a decrease in the rate of efferocytosis will result in the same steady increase in PI+ neurons. The number of PI+ neurons will grow larger the longer the imbalance in the rates of apoptosis vs. efferocytosis persists.

**Fig. 5.**
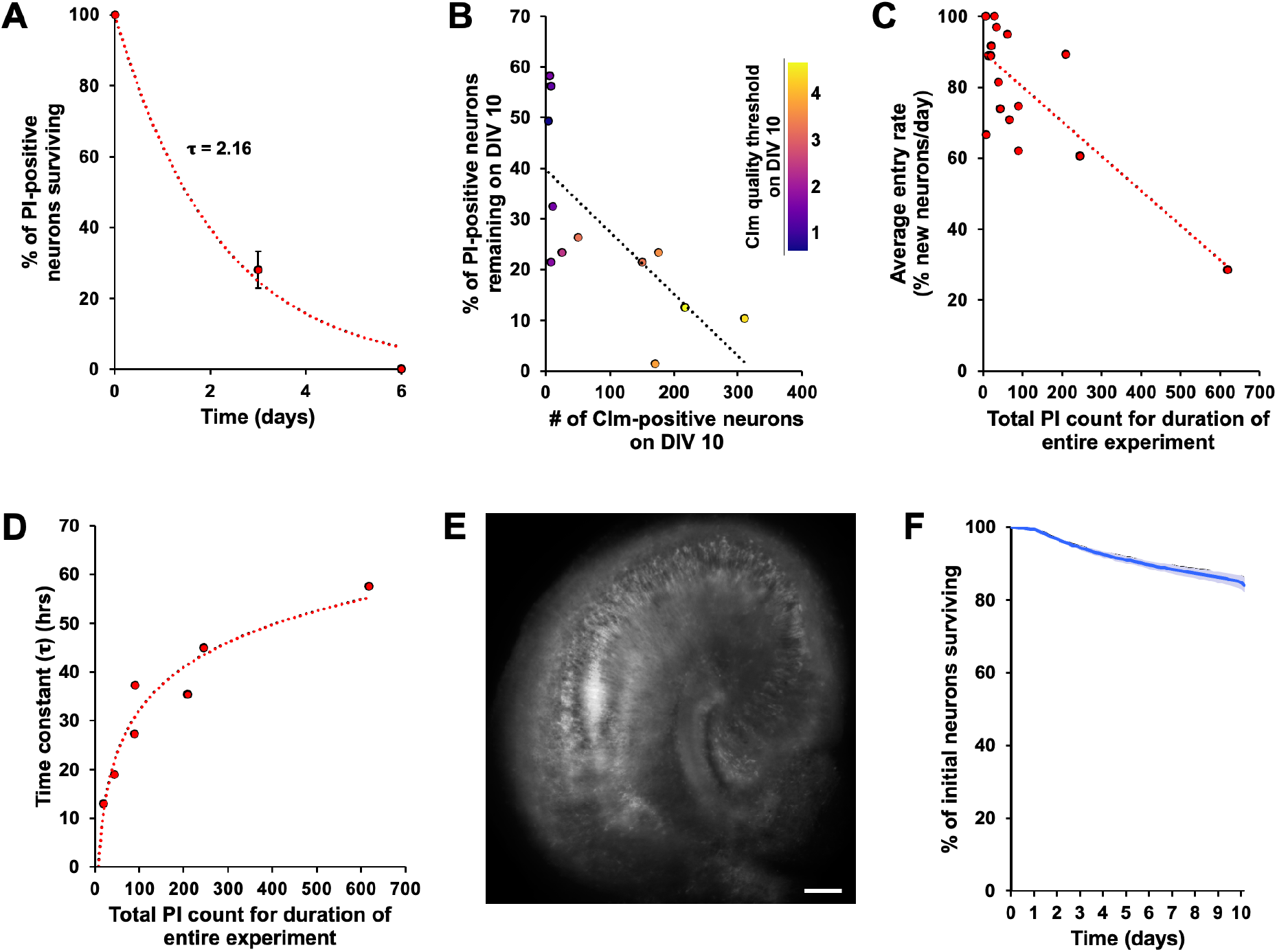
Survival of propidium iodide-positive and fluorescent protein-positive neurons. **(A)** Propidium iodide-positive neurons can survive for days. Organotypic hippocampal slices that expressed the fluorescent protein Clm were given a single dose of the nuclear cell death indicator propidium iodide on DIV 7, then imaged, and subsequently imaged again on DIVs 10 and 13. PI-positive cells were counted using the ImageJ plugin TrackMate. The initial PI count was 967 ± 106.1, and the time constant of the progressive loss of PI-positive cells was 2.16 days (n = 12 slices). **(B)** Propidium iodide-positive neurons survive for longer in less healthy slices. In the same experiment as in (A) there was a correlation between the percentage of the initial PI-positive population that remained on DIV 10, and the number of Clm-positive neurons present on DIV 10 (R-squared = 0.5123, p = 0089, n = 12). TrackMate’s Quality threshold for the Clm-positive cells in each slice (see color bar) also correlated with these measurements. **(C)** The entry rate into the PI-receptive pool does not increase in slices with higher PI counts. Slices were imaged for 72 hrs and TrackMate was used to identify new PI-positive neurons on each day. For each slice the percentage of PI-positive neurons that was newly PI-positive on each day was found, and then the average of all days for that slice was taken. This average was then plotted against the total number of PI-positive neurons counted over the entire 72 hr period. n = 17 slices. **(D)** Propidium iodide-positive neurons survive for longer in slices with higher PI counts. Slices were imaged for 72 hrs and TrackMate was used to track the initial PI-positive neuron population. The time constant was determined from the progressive loss of the initial PI-positive cells for each slice, and plotted against the total number of PI-positive neurons counted over the entire 72 hr period. n = 7 slices. **(E)** Example image of plentiful jRCaMP1a expression in an organotypic slice imaged at DIV 4. Scale bar = 100 μm. **(F)** FP+ neurons can survive for weeks, or longer, during seizures. Organotypic slices infected with AAV-syn-jRCaMP1a and AAV-syn-cre-nls-GFP were imaged every 4 hours from DIV 7 to DIV 17. GFP-positive cells were counted and tracked using ImageJ and TrackMate. After 10 days of imaging 84.03 ± 2.01 % of neurons (7,134 out of 8,486, n = 9 slices) were still visibly fluorescent and morphologically intact, indicating that the vast majority of neurons can survive for a considerable length of time despite the presence of seizures.

To measure the rate at which neurons became PI+, the rate of *entry* into the PI-receptive pool was measured using serial applications of PI to another group of organotypic hippocampal slice cultures at DIV 7 to 10. The number of PI-positive neurons was counted on each day (Supplemental Table 1) and neurons were tracked across multiple days (Supplemental Figure 1). By determining the number of newly-stained PI+ neurons on each day, we found the average daily entry rate of neurons into the PI-receptive pool to be 81% of the extant PI-receptive pool (Figure 5C). Additionally, when the initial PI-positive population in these slices is tracked and the loss of these cells is used to derive the time constants of monoexponential decay, those 𝛕 values are in line with those seen in Figure 5A with the average 𝛕 being 33.52 ± 5.77 hrs, or 1.40 ± 0.24 days (Figure 5D). There was a strong, nonlinear correlation between the time constant for clearance of PI+ neurons and the total number of PI-positive neurons counted over the entire 72 hr period (Figure 5D). This correlation suggests that the ratio of dying neurons to active microglia varies from slice to slice, and that slices with more dying neurons have longer time constants. We conclude that the maximum rate of microglial efferocytosis of dying neurons can be exceeded when too many neurons undergo apoptosis at the same time. This is the condition represented by the system in Figure 4C. Preparations in which efferocytosis rates was saturated, i.e. preparations with high ratios of dying neurons to active microglia, would be expected to have linear rates of PI clearance. However, the development of seizures in the first two weeks in vitro (38, 40) may have slowed efferocytosis and distorted this linearity.

The average rate of entry into the PI+ pool was 81% of the PI+ pool size (Figure 5C). From this it might be concluded that single-time-point staining is a reasonable measure of the rate of apoptosis. However, there was no positive correlation between entry rates and size of the PI+ pool (Figure 5C). Although the pool size variance was large, even if outliers are excluded, there is still no correlation between PI pool size and the rate of entry into the PI+ pool. We had previously assumed (38) that the rate of entry into the PI+ pool would be the primary determinant of the size of the PI+ pool: i.e., in single-time-point analyses, the number of PI+ neurons is a biomarker for the rate of neuronal death. But this is not the case, at least in the organotypic slice preparation (Figure 5C). Rather, the rate of exit from the pool is the primary determinant of the size of the pool of PI+ neurons. This is because PI-positive cells remain in the pool longer when the size of the pool itself is larger (Figure 5D).

The PI results demonstrate wide variation in the rates of exit from the PI receptive pool, and the corresponding changes in pool size (Figure 5D). This is schematized by the microglial axis of Figure 1. The rates of entry into the PI+ pool appear to be the best measure of ongoing cell death, because these neurons have most recently committed to apoptosis. However even the rate of entry into the PI+ pools comes with a caveat, because the time between commitment to apoptosis and entry into the PI-receptive pool is not known. It is possible that all these new PI+ cells committed to apoptosis at the time of slice preparation, or it may be that the newly PI+ cells represent only newly apoptotic cells that had survived slicing unscathed but subsequently committed to apoptosis, or a combination of these two populations.

We therefore used loss of fluorescent protein emission as another assay for neuronal death in serially imaged hippocampal organotypic slice cultures (41–43, 54, 55). Slice cultures infected with AAV-syn-jRCaMP1a and AAV-syn-cre-nls-GFP were imaged every 4 hours from DIV 7 to DIV 17 using a single-photon microscope with a robotic 6-well stage built into a tissue culture incubator (58). GFP-positive neurons were counted and tracked using ImageJ and TrackMate. Spontaneous recurrent seizure activity was present in all slices, assayed by jRCaMP activity (Figure 5E) (40, 58). By DIV 17, 84.03 ± 2.01 % of neurons (7,134 out of 8,486, n = 9 slices) were still visibly fluorescent (Figure 5F). Thus longitudinal assessment of neuronal viability by fluorescence quenching indicated a low rate of neuronal death, about 1% of the healthy fluorescent pool per day, despite the ongoing presence of seizures.

Ideally, we could compare the rates of death from fluorescence extinction directly to the rate of death measured by newly PI+ neurons. However, comparing the newly PI+ rate to the number of FP+ neurons directly is not possible because we do not know the fraction of healthy neurons that are FP+ in these experiments. Our recent published experience with this preparation is that 23% of neurons whose health was established by NeuN and phosphoS6 staining are FP+ after AAV-mediated expression (40). Thus, if 1% of healthy cells enter the apoptosis pathway every day, but only 1 in 4 healthy cells are FP+, then we would expect the total number of neurons entering the apoptototic pathway to be 4 times 1%, i.e. 4% of FP+ neurons. Using Figure 2C, 4% of 3,488 FP+ neurons in 12 slices is 140 neurons entering the apoptotic pathway per day, or 12 neurons per slice per day. Yet if 81% of the PI+ pool of 23,697 PI+ neurons become newly PI+ each day, then 19,195 neurons per day became newly PI+ in 12 slices, so that the newly PI+ rate per day per slice is 1,600. Of course one must allow for variability in slice health, fluorescent protein expression, and PI staining, as well as the fact that the number of PI+ cells is strongly negatively correlated to time post-slicing (38) and so this number will not remain high for the entire lifetime of a slice. But even allowing for this variability, this mismatch between the low rate of apoptosis initiation (12 neurons per slice per day) and the higher rate of newly PI+ measures of neuronal death (1,600 neurons per slice per day) indicates that there is a large pool of FP-neurons that are not healthy (i.e. would not express fluorescent protein) but not yet PI+. These neurons may have entered the apoptotic pathway, but not yet become PI+. We therefore examined this phase of apoptosis.

### AM dye uptake relative to other biomarkers of apoptosis

To address the interval between the time that neurons undergoing apoptosis lose fluorescence emission and the time they become PI+, we exploited a serendipitous finding that in organotypic slices, organic acetoxymethyl ester (AM) dyes selectively stain neurons undergoing apoptosis. AM dyes are esterified to an acetoxymethyl moiety to enhance membrane permeability. Hydrolysis of the ester bond by intracellular esterases renders the dye fluorescent as well as less membrane permeable, “trapping” the fluorescent dye in the neuronal cytoplasm (59). This technique was originally described for cells in suspension. Techniques incorporating pressure, solvents, and detergents have been described to load neurons and astrocytes in situ with AM dyes (60, 61). We found that in hippocampal organotypic slice cultures, overnight incubation of slices with AM dyes selectively stained neurons undergoing apotosis (see Methods). This staining pattern after overnight incubation was consistently observed with both sodium-binding benzofuran isophthalate acetoxymethyl ester (SBFI-AM) and FuraRed-AM across a range of slice conditions and experimental protocols.

First, we confirmed that AM dye-positive cells were neurons by infecting slices with glial fibrillary acidic protein (GFAP)-based virus GFAP-eGFP, which is exclusively expressed in astrocytes. SBFI-AM and GFP-positive cell populations were clearly distinct (Supplemental Figure 2), confirming that neurons load preferentially with AM dyes with the protocols we used in the organotypic hippocampal slice preparation.

Next, we determined that FP-positive neurons quench *in vivo* after acute hypoxic-ischemic injury. Acute ischemic stroke in anesthetized Clm mice induced by Rose Bengal photothrombosis (62–64) resulted in widespread fluorescent protein quenching (Supplemental Figure 3). CFP and YFP emission intensities decreased to 45.26 ± 4.53 % and 51.64 ± 2.98 % of their initial values respectively after 75 min (n = 12 neurons from 3 mice), whereas in a control experiment where photothrombosis was not performed CFP and YFP remained at 97.20 ± 1.61% and 87.10 ± 1.70 % of their initial values respectively after 90 min (n = 12 neurons from 1 mouse) (Figure 6A). This affirmed the utility of fluorescent protein quenching as a biomarker for neuronal death after acute as well as degenerative (41–43, 54, 55) injury.

**Fig. 6.**
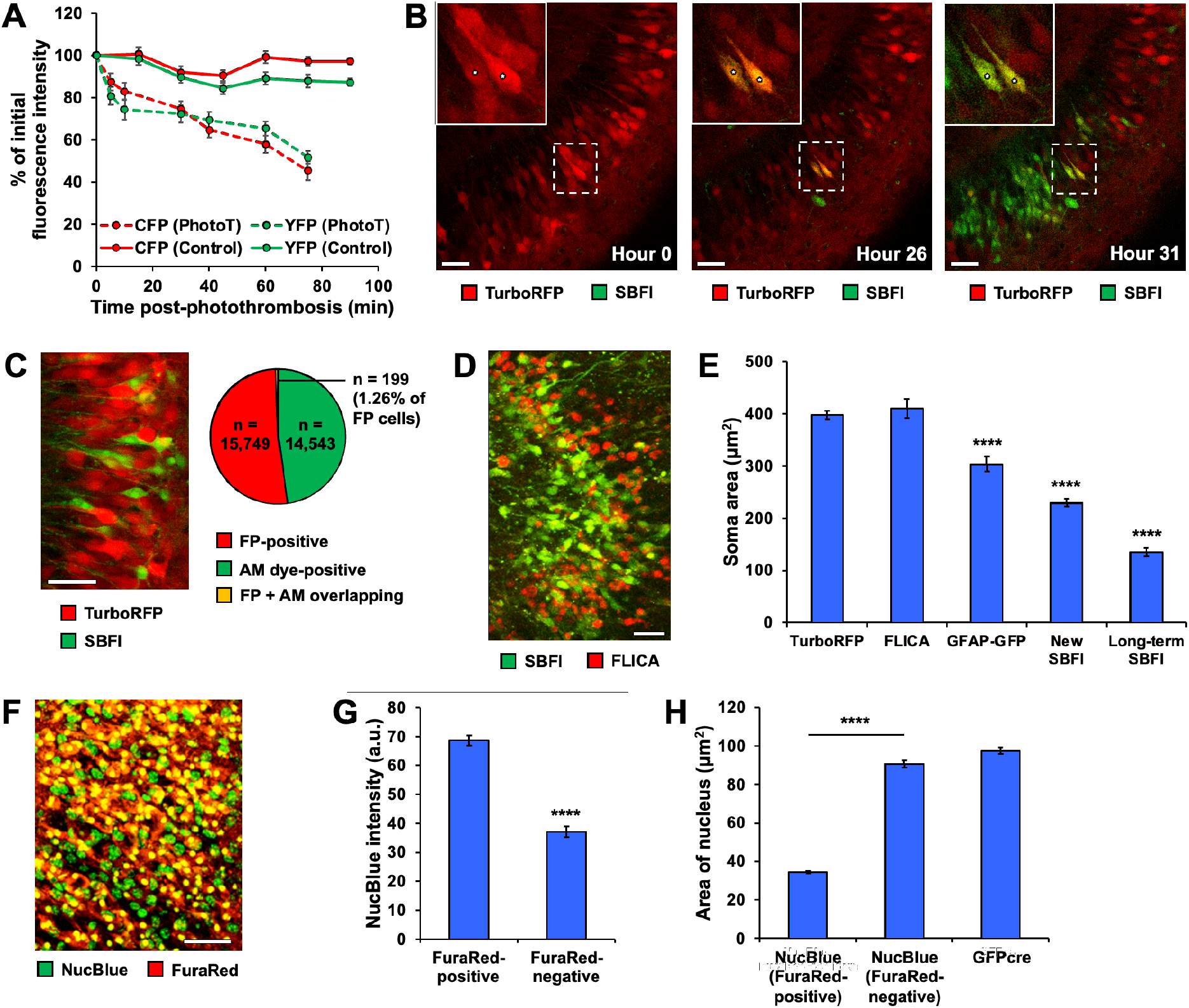
AM dye-positive neurons are undergoing apoptosis. **(A)** Quantification of CFP and YFP quenching *in vivo* post-photothrombosis. Photothrombosis *in vivo* reduced CFP and YFP emission intensities to 45.26 ± 4.53 % and 51.64 ± 2.98 % of their initial values respectively after 75 min (n = 12 neurons from 3 mice), whereas in a control experiment where photothrombosis was not performed CFP and YFP remained at 97.20 ± 1.61% and 87.10 ± 1.70 % of their initial values respectively after 90 min (n = 12 neurons from 1 mouse). **(B)** Quenching of fluorescent proteins and uptake of AM dyes occur concurrently. Depriving an organotypic slice of fresh culture media beyond the normal media change interval caused widespread quenching of TurboRFP over a ∼24 hour period, at which point many neurons also began to take up SBFI-AM. Inset: example neurons (indicated with “*”) that began as TurboRFP-positive and SBFI-negative, briefly became positive for both, and then rapidly became SBFI-positive and TurboRFP-negative. Scale bars = 50 μm. **(C)** AM dye-positive neurons do not express fluorescent proteins. Example image showing that SBFI-positive neurons and TurboRFP-positive neurons are two separate populations (left). Quantification of this lack of overlap, showing that across 14,543 SBFI-positive neurons and 15,749 neurons positive for either TurboRFP or Clm, only 199 of the FP-positive neurons (or 1.26%) also exhibited SBFI-positivity (right). n = 22 SBFI + TurboRFP slices, n = 3 SBFI + Clm slices. Scale bar = 50 μm. **(D)** AM dye-positive neurons are largely FLICA-negative. Neurons that take up SBFI-AM rarely stained positive for FLICA. Slice imaged on DIV 7. Scale bar = 25 μm. **(E)** AM dye-positive neurons exhibit progressive cell shrinkage. FLICA-positive neurons had the same somatic area as TurboRFP-positive neurons (410.06 ± 18.37 μm^2^ vs. 397.56 ± 8.31 μm^2^, n = 80 and 80, p = 0.5361). GFAP-GFP-positive astrocytes had smaller soma than TurboRFP-positive neurons (303.75 ± 14.43 μm^2^ vs. 397.56 ± 8.31 μm^2^, n = 80 and 80, p < 0.0001). Neurons that had been SBFI-positive for less than 24 hours had a reduced somatic area compared to TurboRFP neurons (229.72 ± 7.14 μm^2^ vs. 397.56 ± 8.31 μm^2^, n = 80 and 80, p < 0.0001), and neurons that had been SBFI-positive for longer than 24 hours had an even smaller somatic area (135.28 ± 7.72 μm^2^ vs.397.56 ± 8.31 μm^2^, n = 60 and 80, p < 0.0001). **(F)** AM dye-positive neurons exhibit distinctive chromatin morphology. Example image showing that the nuclei of FuraRed-positive neurons, stained with NucBlue, are visually distinct from the nuclei of FuraRed-negative neurons. Scale bar = 50 μm. **(G)** AM-dye positive neurons exhibit chromatin condensation. Quantification of NucBlue staining shown in (F), wherein FuraRed-positive neurons had greater NucBlue intensity than FuraRed-negative neurons (68.61 ± 1.67 vs. 37.04 ± 1.80, n = 25 vs. 25, p < 0.0001). **(H)** Nuclear area of neurons staining with FuraRed AM is smaller than both FuraRed-negative neurons (34.51 ± 0.59 μm^2^ vs. 90.70 ± 1.84 μm^2^, n = 157 vs. 157, p < 0.0001) and neurons whose nuclei expressed GFPcre (34.51 ± 0.59 μm^2^ vs. 97.47 ± 1.60 μm^2^, n = 159 vs. 113, p < 0.0001).

We next determined the extent of AM dye uptake over the course of fluorescent protein quenching in organotypic hippocampal slice cultures. Healthy TurboRFP-positive neurons exhibited no SBFI-AM staining (Figure 6B). However, if the same neurons were induced to initiate apoptosis by media starvation (slice cultures kept in the same media for 24 hours beyond the typical 3-day interval for exchanging spent for fresh media), TurboRFP fluorescence began to quench and scattered neurons began to take up SBFI (Figure 6B middle). After just 5 additional hours, many neurons now stained with SBFI-AM (Figure 6B right). Additionally, individual neurons could be observed quenching TurboRFP and taking up SBFI-AM simultaneously, and thus visibly transitioning from red to yellow to green (Figure 6B insets). This indicates that fluorescent protein quenching and an increased propensity to load with cell permeant AM dyes are concurrent processes. At this early stage, SBFI-AM+ neurons retained their dendritic processes (Figure 6B).

In control slices not deprived of fresh media, healthy neurons that expressed robust emission from transgenically expressed fluorescent proteins TurboRFP or Clm only rarely stained with SBFI-AM: only 199 FP-positive neurons (or 1.26%) exhibited both SBFI-AM and TurboRFP/Clm simultaneously, whereas 15,749 neurons were observed that were FP-positive but SBFI-AM negative, and 14,543 neurons were observed that were SBFI-positive but FP-negative (n = 22 SBFI + TurboRFP slices, n = 3 SBFI + Clm slices) (Figure 6C). This small amount of overlap indicates that the AM dye-positive population is comprised of neurons whose fluorescent proteins have quenched at the onset of apoptosis.

SBFI-AM positive neurons were rarely FLICA-positive (Figure 6D). This suggests that increased caspase activity is not only an early step in neuronal apoptosis (as seen in Figure 3A), but also a transient one, else SBFI-AM positive neurons would continue to be FLICA-positive throughout their remaining lifetime. The lack of FLICA staining of SBFI-AM positive neurons also indicated that SBFI-AM is not toxic under these conditions, i.e. it was not inducing healthy neurons to initiate apoptosis and thereby become both SBFI- and FLICA-positive (65).

### Morphological evidence of apoptosis in AM dye-positive neurons

Programmed cell death is accompanied by somatic shrinkage (66). AM dye-positive neurons exhibited progressive cell shrinkage during serial imaging experiments (Figure 6E), with neurons that had been SBFI-AM positive for less than 24 hours having a reduced somatic area compared to TurboRFP-positive neurons (229.72 ± 7.14 μm^2^ vs. 397.56 ± 8.31 μm^2^, n = 80 and 80 neurons, p < 0.0001). Neurons that had been SBFI-AM positive for longer than 24 hours had an even smaller somatic area (135.28 ± 7.72 μm^2^ vs. 397.56 ± 8.31 μm^2^, n = 60 and 80 neurons, p < 0.0001). FLICA-positive neurons had the same somatic area as TurboRFP-positive neurons (410.06 ± 18.37 μm^2^ vs. 397.56 ± 8.31 μm^2^, n = 80 and 80, p = 0.5361), whereas GFAP-GFP-positive astrocytes had smaller soma than TurboRFP-positive neurons (303.75 ± 14.43 μm^2^ vs. 397.56 ± 8.31 μm^2^, n = 80 and 80, p < 0.0001). This supports the idea that most FLICA-positive cells are neurons, since their area matched that of TurboRFP-positive neurons so closely. However, the distribution of somatic area values for FLICA-positive cells was broader than that of TurboRFP-positive neurons (Supplemental Figure 4). This could mean that some FLICA- positive cells were, for example, astrocytes. However, the vast majority of cells undergoing apoptosis after acute brain injury are neurons (67, 68), and neurons are also known to be much more vulnerable than astrocytes following oxygen-glucose deprivation (69). Thus, it is more likely in our case that some FLICA-positive neurons had already started to shrink, and as a result they skewed closer to the area values of newly SBFI-positive cells.

Chromatin condensation is another morphological marker of programmed cell death (66). The apoptotic status of AM dye-positive neurons was further confirmed via NucBlue staining. After overnight incubation of slices loaded with FuraRed-AM with 4 drops of NucBlue, the nuclei of FuraRed-AM positive neurons appeared markedly smaller and stained more intensely with NucBlue than did FuraRed-negative neurons (Figure 6F). This strongly suggests that AM dye-positive neurons are undergoing chromatin condensation, another hallmark of apoptosis (29). This was quantified by comparing both the NucBlue emission intensity and the area of the nuclei. The nuclei of FuraRed-positive neurons had significantly greater NucBlue emission than FuraRed-negative neurons (68.61 ± 1.67 a.u. vs. 37.04 ± 1.80 a.u., n = 25 vs. 25, p < 0.0001) (Figure 6G), and also had significantly smaller nuclear area (34.51 ± 0.59 μm^2^ vs. 90.70 ± 1.84 μm^2^, n = 157 vs. 157, p < 0.0001) (Figure 6H). As an additional confirmation, the nuclear area of neurons positive for both FuraRed and NucBlue were compared to those of neurons expressing nuclear GFP due to infection with AAV9.hSyn.HI.eGFP-Cre.WPRE.SV40. The FuraRed-positive neurons were again found to have a smaller nuclear area than the healthier GFPcre-positive neurons (34.51 ± 0.59 μm^2^ vs. 97.47 ± 1.60 μm^2^, n = 159 vs. 113, p < 0.0001) (Figure 6H). Based on these morphological features we can also conclude that these neurons are not undergoing ferroptosis. Ferroptosis is another form of programmed cell death characterized by lipid peroxidation and the presence of iron. However, ferroptosis does not involve cell shrinkage or chromatin condensation (70–72).

We compared the time course of AM dye-staining and quenching of the emission of fluorescent proteins as indicators of neuronal death. As in Figure 6B, depriving slices of fresh media past when they would normally be fed resulted in widespread AM dye uptake and fluorescent protein quenching. 15 neurons from 5 such slices were selected randomly from the pool of neurons which underwent the FP-to-AM transition and could be visualized and tracked for the duration of the experiment. Using ImageJ, SBFI-AM and TurboRFP fluorescence intensity were both quantified (Supplemental Figure 5A-C). The ratio of SBFI-AM to TurboRFP intensity for a given neuron was calculated, and the time interval at which this ratio exhibited its largest increase was taken to be time = 0 for the observable initiation of cell death. All other time points for that neuron were aligned accordingly. The largest drop in TurboRFP fluorescence (90.28 ± 3.84 % of maximum to 58.78 ± 5.85 %) and the largest increase in SBFI fluorescence (12.95 ± 6.22 % of maximum to 51.08 ± 5.83 %) were found to occur during the same time interval (Supplemental Figure 5A). Graphing the SBFI/TurboRFP ratio itself over time further highlights how synchronous the two processes are, with the ratio experiencing its largest increase (0.13 ±0.05 to 1.13 ± 0.13) at t = 0 (Supplemental Figure 5B). Shrinkage of the soma, another indicator of apoptosis, aligned very closely with these metrics, with largest drop in 2-D soma area (90.07 ±5.08 % of maximum to 70.03 ± 3.06 %) occurring during the same time interval as did maximum SBFI uptake and maximum TurboRFP quenching (Supplemental Figure 5C). This suggests that all three processes (SBFI staining, fluorescent protein quenching, and cell shrinkage) occur concurrently as apoptosis begins. This is visualized in Supplemental Figure 5D, where a slice with plentiful TurboRFP expression and sparse SBFI uptake at Hour 0 is contrasted with the same slice at Hour 96, where 89% of TurboRFP-positive neurons have quenched and SBFI uptake is pervasive. Thus, AM dye uptake can be a useful complement to enhance the temporal resolution of fluorescent protein quenching as an assay for cell death after acute injury.

Figure 6 and Supplemental Figure 5 establish that AM dye staining is an early biomarker of apoptosis in the hippocampal organotypic slice preparation. To use AM dyes as a biomarker of apoptosis, it is necessary to know how well the biomarker performs throughout the entire process of apoptosis. Neuronal apoptosis ends in microglial efferocytosis (3, 19, 20). To test whether neurons undergoing apoptosis remained AM-dye positive from the onset of apoptosis through the final stage of efferocytosis, microglial efferocytosis of neurons was evaluated in the organotypic hippocampal slice preparation. Isolectin GS-IB4 conjugated to Alexa Fluor 594 was used to visualize activated microglia. Slices were incubated with 10 μg/mL of isolectin for a minimum of 150 minutes, and were incubated overnight with 27 μM SBFI.

### Survival duration of AM dye-positive neurons

Isolectin-positive microglia were frequently observed in close proximity to, and in direct contact with, SBFI-positive cells (Supplemental Figure 6A). When imaged over several days, the entire timeline of the microglial-neuronal interaction could be observed, wherein an SBFI-AM positive cell is completely engulfed and consumed by microglia (Supplemental Figure 6B). Perhaps owing to the large number of apoptotic neurons waiting to be efferocytosed relative to the number of microglia available to consume those neurons, the duration of microglial efferocytosis was highly variable. 2 slices were observed via serial imaging and 18 SBFI-positive neurons were selected randomly from the pool of neurons which could be observed undergoing the entire engulfment process. The amount of time those neurons spent in contact with microglia before being fully engulfed (“Contact” phase), the time spent being engulfed before the neurons underwent terminal cell shrinkage (“Engulfment” phase”), and the time spent undergoing terminal neuronal shrinkage (“Shrinkage” phase) were all quantified (Supplemental Figure 6C). The Contact phase lasted an average of 10.17 ± 4.71 hours, with the longest observed Contact taking 37 hours and the shortest taking only 1 hour. The Engulfment phase lasted an average of 16.43 ± 5.74 hours, with the longest Engulfment taking 55.5 hours and the shortest taking 1 hour. Lastly, the Shrinkage phase lasted an average of 27.08 ± 5.19 hours, with the longest taking 59 hours and the shortest taking 4.5 hours. For the neurons which could be observed undergoing all three phases, the average total duration of the entire engulfment process was 46.06 ± 7.86 hours (n = 8). This, of course, does not take into account the additional time required for microglia to find their target neurons and move into contact with them. This would be expected to take longer in regions that are dense with dying neurons, and thus would be a source of additional variability. Notably, in a slice with TurboRFP-positive neurons that was imaged over 3 days, no instances of microglial engulfment of these healthy neurons could be seen, despite the number of microglia present in the area increasing over the course of the experiment (Supplemental Figure 7). This further supports the idea that microglia selectively target apoptotic neurons.

These data indicate that AM dyes permeate neurons undergoing apoptosis from the time of fluorescence quenching (Figure 6B) through terminal efferocytosis (Supplemental Figure 6). AM dyes therefore can be used to estimate the time course of neuronal apoptosis after acute, widespread neuronal injury in the organotypic slice preparation. Slices expressing TurboRFP and incubated with SBFI in standard culture conditions were imaged every 3-5 days for an 18-day period (Figure 7A). A subset of SBFI-AM positive neurons was manually selected based on the following criteria: neurons that (a) were newly SBFI-AM positive; (b) underwent terminal cell shrinkage and died over the course of the experiment; and (c) were located in regions with sparse SBFI staining, to aid in their precise visual identification over time. These criteria yielded 41 neurons from 4 slices, which were manually tracked for the duration of the experiment. The continued presence of TurboRFP-positive neurons was used to verify that the slices remained healthy overall, and indeed TurboRFP-positivity persisted even as the SBFI-AM positive population dwindled (Figure 7B). The SBFI-positive neurons were found to have a 𝛕 of 4.69 days and a half-life of 3.66 days, with some neurons surviving for up to 12 days (Figure 7C). This data indicates that neurons undergoing apoptosis can persist for up to 12 days as they await microglial efferocytosis, accounting for the delay referred to in the term Delayed Neuronal Death (5). In a separate experiment, slices were loaded with isolectin and SBFI-AM and imaged serially over 5 days. 𝛕 was determined from the loss of the initial SBFI-AM positive cells for each slice, and 𝛕 was then plotted against the ratio of the ratio of the number of isolectin-positive microglia to SBFI-positive neurons, averaged over the first 24 hrs of the experiment (Figure 7D). The negative correlation indicates that in slices with plentiful microglia relative to the number of dying neurons, the microglia can clear those neurons more quickly. This idea is also supported by the data in Figure 5D, where the rate of loss of PI+ neurons decreased with the number of PI+ neurons in the preparation (Figure 5D).

**Fig. 7.**
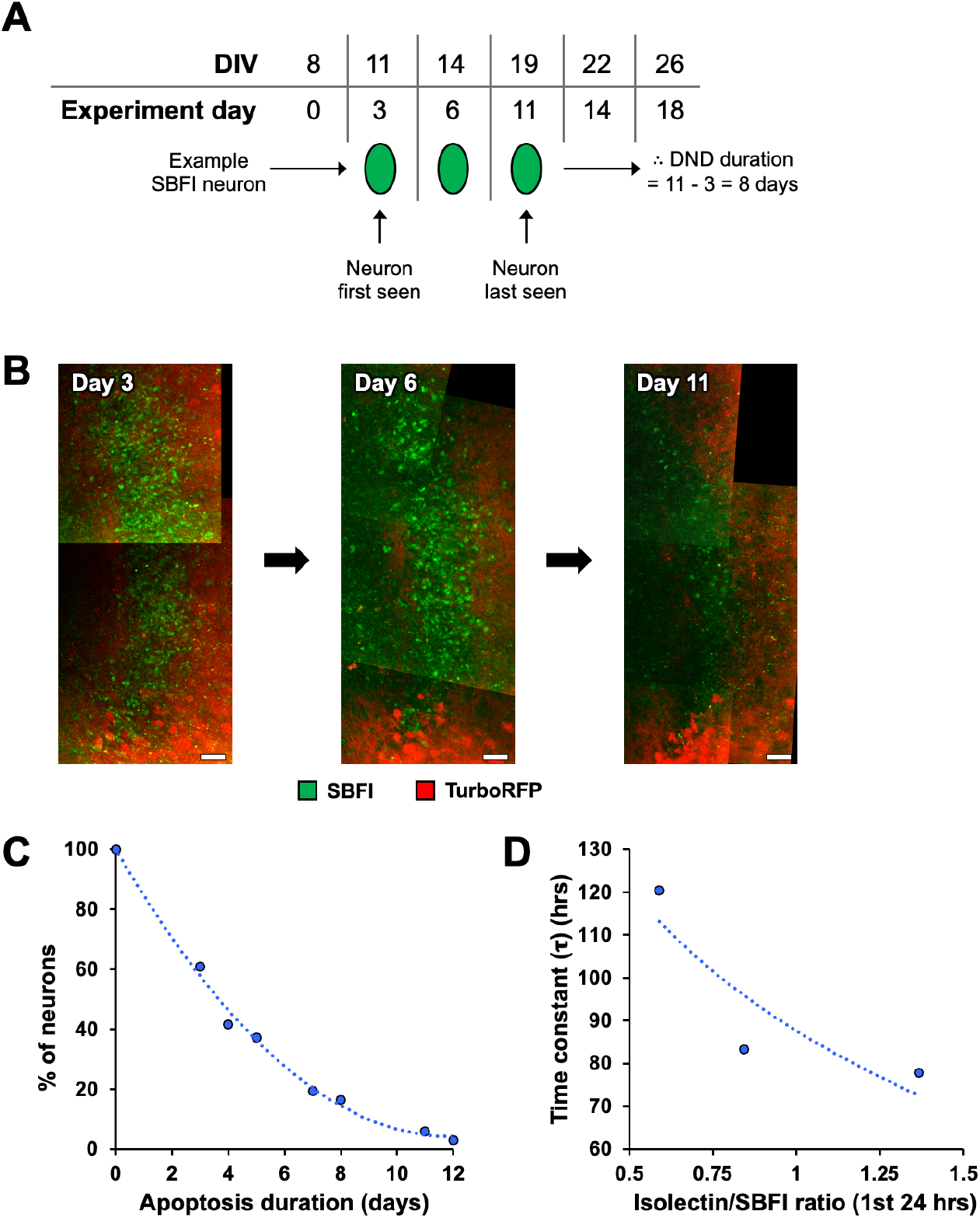
Survival of AM dye-positive neurons. **(A)** Schematic of serial imaging experiment to measure the survival of neurons which take up the cell-permeant AM dye SBFI. Slices were imaged every 3-5 days beginning on DIV 8. The duration of survival of SBFI+ neurons was calculated by subtracting the experiment day on which an SBFI-positive was first seen, from the experiment day on which it was last seen. **(B)** Images of SBFI- and TurboRFP-positive neurons at three time points during an experiment in (B). While TurboRFP-positive neurons persist, SBFI-positive neurons are largely gone by Day 11 of the experiment. Experiment Day 3 = DIV 11. Scale bar = 50 μm. **(C)** SBFI+ neurons can survive for a substantial period of time, consistent with earlier findings indicating that microglia efferocytosis is rate-limiting after widespread neuronal injury. Neurons that became SBFI-positive and died during an 18 day-long experiment had a 𝛕 of 4.69 days and a half-life of 3.66 days, though some were still alive after 12 days. n = 41 cells from 4 slices. **(D)** Survival of SBFI-positive neurons correlates with the relative levels of microglia to their phagocytic targets. In slices stained with isolectin and SBFI, both isolectin-positive microglia and SBFI-positive neurons were imaged over 5 days then tracked with TrackMate. The time constant was determined from the progressive loss of the initial SBFI-positive cells for each slice. 𝛕 was then plotted against the ratio of the number of isolectin-positive microglia to SBFI-positive neurons, averaged over the first 24 hrs of the experiment.

These data demonstrate that AM dyes are biomarkers for neurons throughout apoptosis in hippocampal organotypic slice preparation, from the earliest stages of fluorescence quenching (Figure 6B) to terminal effertocytosis by microglia (Supplemental Figure 6). Biomarkers based on more polar organic dyes such as PI are only positive at late stages of apoptosis (Figure 2), suggesting that membrane permeability must increase during the apoptotic process. Although the dependence of biomarker staining on pathological levels of membrane permeability exploits this unique feature of apoptosis (34), it also raises the possibility of false positive stains. This is because membrane permeabilization, effected by detergents and alcohols (73, 74), is a key step in cytochemical and immunohistochemical staining that allows the biomarkers access to the cytoplasmic and nuclear contents. To be able to evaluate potential false positive biomarker staining, we carefully characterized the membrane permeability of neurons undergoing apoptosis.

### Membrane permeability during neuronal apoptosis after perinatal acute brain injury

Annexin V (Alexa Fluor 594 conjugate) is a biomarker of membrane deterioration during apoptosis. Annexin V selectively stains apoptotic cells by binding to phosphatidylserine, a component of the inner leaflet of the cytoplasmic membrane which is found on the outer leaflet of the membranes during apoptosis (30, 31, 75). SBFI-AM positivity overlapped significantly with positivity for Annexin V. In slices incubated with 50 μL Annexin V per mL of culture media for a minimum of 150 minutes, 491 out of 506 SBFI-positive neurons (or 97.04%) were found to also be positive for Annexin V (Figure 8A).

**Fig. 8.**
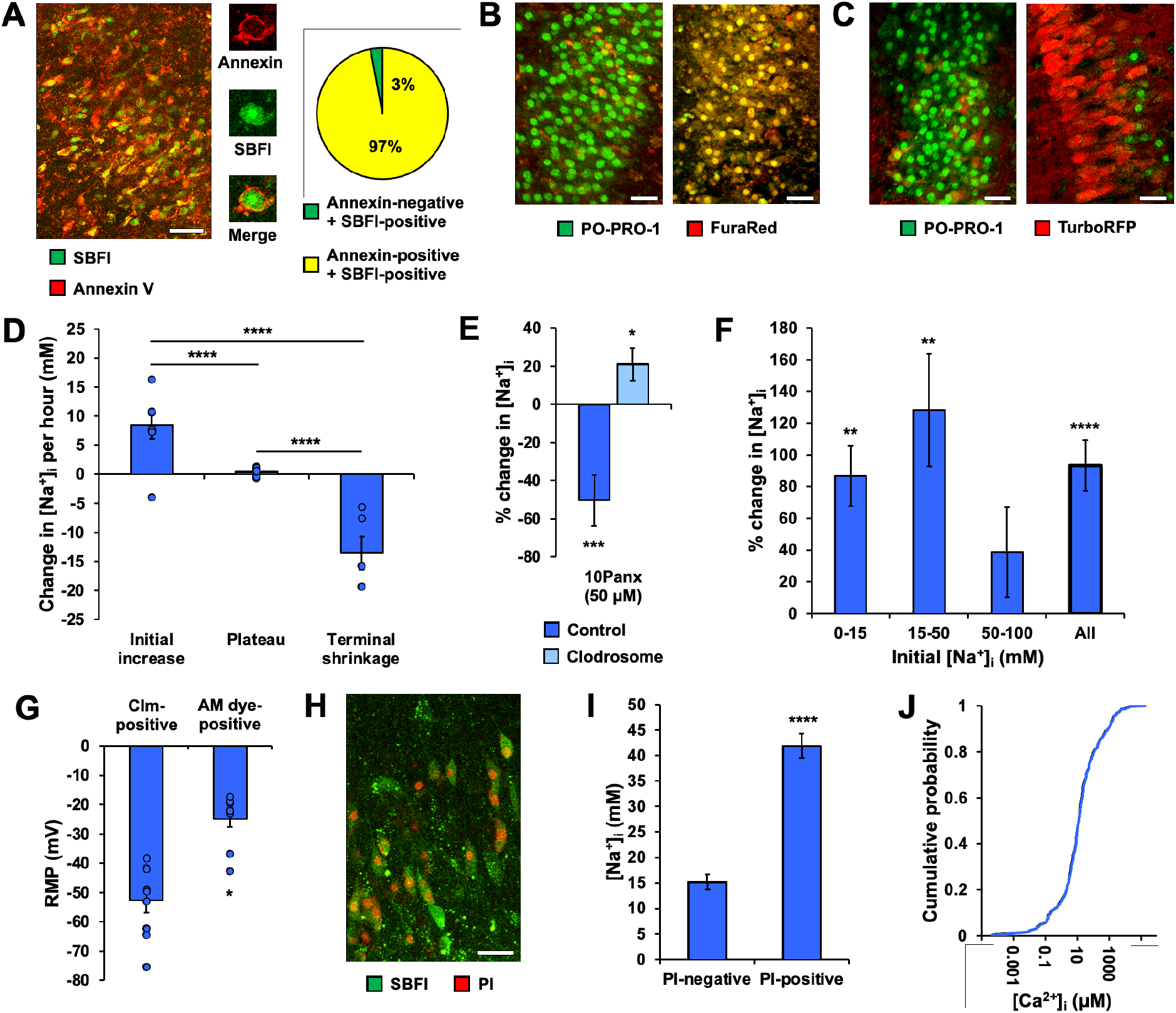
Apoptotic neurons undergo progressive membrane deterioration. **(A)** Evidence of membrane injury in AM dye-positive neurons. Example image showing broad overlap between SBFI-positivity and membrane injury indicated by Annexin V staining of phosphatidylserine moieties in the cytoplasmic membrane (left). Larger example images of an individual neuron that is positive for both SBFI and Annexin V (middle). Quantification of SBFI/Annexin V overlap, showing that 97% of SBFI-positive neurons are also positive for Annexin V (right). n = 506 cells from 1 slice. Scale bar = 50 μm. **(B)** AM dye-positive neurons exhibit extensive Pannexin pore expression. Example images showing a group of neurons with PO-PRO-1-positivity (left), and another group in the same slice with extensive FuraRed- positivity where nearly all FuraRed neurons are also PO-PRO-1-positive (right). Scale bars = 25 μm. **(C)** FP-positive neurons do not exhibit Pannexin pore expression. Examples images showing a group of neurons with PO-PRO-1-positivity (left), and another group in the same slice with TurboRFP-positivity that are all PO-PRO-1-negative (right). Scale bars = 25 μm. **(D)** Apoptotic neurons accumulate intracellular sodium ions. In the 5 hours following SBFI uptake [Na^+^]_i_ increased by 8.51 ± 2.35 mM/hour. n = 7 cells from 2 slices. Later in the apoptosis process, [Na^+^]_i_ largely plateaued, only increasing by 0.44 ± 0.08 mM/hour over 24 hours. n = 46 cells from 4 slices. Finally, during terminal cell shrinkage [Na^+^]_i_ rapidly decreased by 13.53 ± 2.93 mM/hour in the 3 hours before cell death. n = 5 cells from 5 slices. p < 0.0001 for Initial increase vs. Plateau. p < 0.0001 for Plateau vs. Terminal cell shrinkage. p < 0.0001 for Initial increase vs. Terminal cell shrinkage. **(E)** Blocking pannexin-1 hemichannels with 50 μM of the 10Panx for 30 min decreased [Na^+^]_i_ (−50.28 ± 13.49 %, p = 0.0004, n = 64 neurons from 5 slices), whereas 10Panx application in microglia-depleted slices increased [Na^+^]_i_ (21.04 ± 8.55 %, p = 0.0162, n = 74 neurons from 3 slices). **(F)** Blockage of Na/K ATPases increases [Na^+^]_i_ in apoptotic neurons. Acute application of 10 μM ouabain for 30 min increased [Na^+^]_i_ by 86.84% in neurons with an initial [Na^+^]_i_ between 0-15 mM (n = 84 cells, p = 0.0048), 128.19% in neurons with an initial [Na^+^]_i_ between 15-50 mM (n = 25 cells, p = 0.0017), had no effect on neurons with an initial [Na^+^]_i_ between 50-100 mM (n = 6 cells, p = 0.2692), and with all cells combined increased [Na^+^]_i_ by 93.32% (n = 115 cells, p < 0.0001). n = 6 slices. **(G)** Apoptotic neurons have depolarized resting membrane potentials. FuraRed-positive neurons had RMPs that were highly depolarized compared to Clm-positive neurons (−24.78 ± 2.93 mV vs. −52.78 ± 4.13 mV, n = 9 cells for each, p < 0.0001). **(H)** Apoptotic neurons have compromised nuclear membranes. An example image showing that many SBFI-positive neurons become PI-positive. Scale bar = 25 μm. **(I)** Compromised nuclear membranes are a feature of late-stage apoptosis. SBFI-positive neurons that are also PI-positive had higher [Na^+^]_i_ than those that are PI-negative (41.93 ± 2.35 mM vs. 15.21 ± 1.47 mM, n = 106 cells vs. 221 cells, n = 6 slices, p < 0.0001), indicating that PI-positivity does not occur until relatively late in the apoptosis process. **(J)** Apoptotic neurons demonstrate a wide range of intracellular calcium concentrations (median = 12.81 μM, n = 335).

In addition to membrane deterioration, active processes may underlie the increase in membrane permeability of apoptotic neurons. Pannexin channels are large-conductance ion channels that are inserted into the neuronal membrane after injury (76). AM dye-positivity also overlapped with positivity for PO-PRO-1, a dye which is preferentially taken up by neurons after pannexin 1 (PANX1) channels have been inserted in the cytoplasmic membrane. After overnight incubation with 28 μM of the AM dye FuraRed, and incubation with 5 μM of PO-PRO-1 for a minimum of 180 minutes, many PO-PRO-1-positive cells could be seen, and neurons that were FuraRed-positive were almost certain to also be PO-PRO-1-positive (Figure 8B). Conversely, even in slices with plentiful PO-PRO-1-positivity, PO-PRO-1 never overlapped with TurboRFP-positive neurons (Figure 8C), providing further evidence that FP-positive healthy neurons and neurons with high membrane permeability are separate populations.

To test whether the timing of biomarker positivity reflects the degree of permeability of the cytoplasmic membrane of neurons undergoing apoptosis, we exploited the shift in wavelengths at which AM dyes absorb light due to changes in local ionic concentrations. We used this property to test whether the timing of biomarker positivity reflects the degree of deterioration of the cytoplasmic membrane’s function of separating extra and intracellular fluids (77). We found that AM dye-positive neurons also accumulate intracellular sodium ions, with the rate of change in the intracellular sodium concentration ([Na^+^]_i_) varying widely depending on the stage of apoptosis. Using the ratiometric properties of SBFI-AM, [Na^+^]_i_ was quantified in neurons over the course of 24-hour serial imaging experiments. Neurons that took up SBFI-AM during the experiment exhibited an [Na^+^]_i_ increase of 8.51 ± 2.35 mM/hour (n = 7 cells from 2 slices) over the first 5 hours that they were SBFI-positive, further confirming that a large increase in membrane permeability is correlated with AM dye uptake. Later in apoptosis [Na^+^]_i_ largely plateaued, only increasing by 0.44 ± 0.08 mM/hour (n = 46 cells from 4 slices) over 24 hours. Finally, during microglial engulfment and terminal cell shrinkage the trend reversed and [Na^+^]_i_ rapidly decreased by 13.53 ± 2.93 mM/hour in the 3 hours before cell death (n = 5 cells from 5 slices; Figure 8D). These rates of change in [Na^+^]_i_ were all significantly different (p < 0.0001 for all comparisons). Thus, neurons undergoing apoptosis experience a large early influx of Na^+^, followed by a potentially lengthy period where [Na^+^]_i_ is elevated but only gradually increasing, and then ultimately [Na^+^]_i_ drops precipitously during terminal cell shrinkage and efferocytosis. This final drop is not currently well understood, but it may be due to the near-total engulfment of the neurons by microglia at this stage (Supplemental Figure 6) cutting the neurons off from the high levels of Na^+^ in the extracellular space.

The increase in [Na^+^]_i_ during apoptosis could be due to increased influx, or decreased extrusion of Na^+^. Blockade of pannexin pores with the 50 μM of the Panx-1 mimetic inhibitory peptide 10Panx for 30 min reduced [Na^+^]_i_ by 50.28 ± 13.49 % in preparations with intact microglial populations (Figure 8E). To test whether microglia might contribute to the role of pannexins in neuronal membrane permeability, slices were depleted of microglia by incubating them with 40 μL of liposomal clodronate (Clodrosome) suspension per mL of culture media as previously described (78). In microglia-depleted slices, 10Panx application no longer reduced [Na^+^]i, and in fact increased [Na^+^]_i_ by 21.04 ± 8.55 %. These findings support the ideas that pannexins contribute to the increase in membrane permeability during neuronal apoptosis, that the increase in permeability contributes to the increase in Nai^+^, and that microglia contribute to pannexin functioning (79).

To address a potential role of decreased Na extrusion in the elevation of [Na^+^]_i_ in apoptotic neurons, we manipulated sodium–potassium adenosine triphosphatase (Na/K ATPase) pharmacologically and by inhibition of ATP production. We found that despite their increased membrane permeability and elevated [Na^+^]_i_, neurons in the apoptotic pathway are still producing and utilizing ATP to power transport mechanisms and regulate ionic gradients. Acute application of 10 μM ouabain (a blocker of Na/K ATPases) for 30 minutes increased [Na^+^]_i_ by 86.84% in neurons with an initial [Na^+^]_i_ between 0-15 mM (n = 84 neurons from 6 slices, p = 0.0048), 128.19% in neurons with an initial [Na^+^]_i_ between 15-50 mM (n = 25 neurons from 6 slices, p = 0.0017), had no effect on neurons with an initial [Na^+^]_i_ between 50-100 mM (n = 6 neurons from 6 slices, p = 0.2692), and with all neurons combined increased [Na^+^]_i_ by 93.32% (n = 115 neurons from 6 slices, p < 0.0001) (Figure 8F). Using whole-cell patch clamp recordings of FuraRed- and Clm-positive neurons from the same slices, the resting membrane potential (RMP) of the FuraRed-positive neurons in culture media (extracellular potassium concentration = 5.3 mM) was found to be greatly depolarized with respect to the RMP of Clm-positive neurons (− 24.78 ± 2.93 mV vs. −52.78 ± 4.13 mV, n = 9 vs. 9, p < 0.0001) (Figure 8G). These results suggest that neurons undergoing apoptosis are utilizing ATP to remove excess Na^+^ from the intracellular space, but that as the membrane permeability continues to increase over time active transport is overwhelmed, resulting in membrane depolarization and increases in intracellular sodium.

The progressive increase in membrane permeability is not limited to the cytoplasmic membranes. Many SBFI-positive neurons also demonstrated nuclear PI staining (Figure 8H), indicating that apoptotic neurons eventually experience a loss of nuclear membrane integrity. Indeed, PI staining would appear to be selective for late-stage apoptotic neurons that have already experienced a significant increase in membrane permeability, as SBFI-positive neurons that were also PI-positive had higher [Na^+^]_i_ than those that were PI-negative (41.93 ± 2.35 mM vs. 15.21 ± 1.47 mM, n = 106 cells vs. 221 cells, n = 6 slices, p < 0.0001) (Figure 8I).

The increase in the ionic permeability of the cytoplasmic membrane permeability in apoptotic neurons was also supported by measurements of cytosolic calcium (Ca^2+^), using ratiometric determination of intracellular calcium concentrations ([Ca^2+^]_i_) obtained with FuraRed-AM (Figure 8J). As with [Na^+^]_i_, apoptotic neurons exhibited a wide range of [Ca^2+^]_i_ values, with a median value of 12.81 μM (n = 335 neurons), which is well above the 50 nM cytoplasmic calcium concentration of healthy hippocampal pyramidal cells (80).

These findings indicate that after acute injury, neuronal apoptosis progresses over days to weeks. Neuronal apoptosis is characterized by membrane deterioration and increasing membrane permeability that drives many of the biomarkers for apoptosis (29, 31, 35). Neuronal apoptosis terminates with microglial efferocytosis (3, 20). As shown in Figure 5, one-time staining for a biomarker is a poor estimate for the rate of neuronal death, i.e. entry into the apoptotic pathway; in the case of PI staining for cell death in organotypic slices, it was off by 2 orders of magnitude when compared to extinction of fluorescent protein emission. However the data also provide evidence for additional potential sources of error in the assessment of neuronal death. First, some experimental manipulations may affect microglia rather than neurons. These manipulations may affect the rate of efferocytosis, as for example depletion of microglia by Clodrosome (78), and as a consequence affect the number of visible (i.e. PI+) apoptotic neurons (Figure 1). The manipulation may also affect interactions between microglia and neurons other than efferocytosis; for example neurons in microglia-depleted slices did not exhibit a pannexin-mediated increase in sodium permeability (Figure 8E). Second, a manipulation that affects the permeability of the membrane of neurons undergoing apoptosis could increase neuronal staining with biomarkers and artifactually change the size of the pool of biomarker-receptive neurons in Figure 4, as schematized in Figure 1. Third, neurons that necrose or are phagocytosed prior to the assay for neuronal death being performed will lead to an underestimate of neuronal injury. In the next section, we consider how effects of interventions on microglia can lead to misinterpretations of the neurotoxic or neuroprotective impact of experimental interventions.

### Altering microglial activity affects the number of visible dying neurons

As schematized in Figure 4, the number of neurons positive for a biomarker of apoptosis depends on the rates of neuronal entry and exit into the biomarker-receptive pool. Because exit from this pool entails microglial efferocytosis, we tested the possibility that altering the rate of microglial efferocytosis will alter the size of the biomarker-receptive pool of apoptotic neurons. 17 slices (DIV 5-7 at the time of imaging) were depleted of microglia treatment with Clodrosome for 3 days prior to imaging (Figure 9A). The number of PI-positive cells in preparations depleted of microglia were counted relative to the number of PI cells in 6 control slices (Figure 9B). In a three-dimensional volume 590×590 μm in area and extending 80 μm along the z-axis from the surface of the slices, Clodrosome-treated slices had significantly more PI-positive cells than control slices (1836.53 ± 206 vs. 936.17 ± 134, p = 0.0208). Similarly, in a z-axis range extending from 80-120 μm deep from the slice surface, Clodrosome-treated slices again had more PI-positive cells than control slices (373.35 ± 72.30 vs. 34.83 ± 6.37, p = 0.0123). The depth dependence reflects the degree of traumatic injury incurred during slice preparation (36). The rate of microglial efferocytosis comprises the rate of exit from the biomarker receptive pool in the scheme shown in Figure 4, so reducing the exit rate increases the number of neurons in the PI-receptive pool (Figure 4C vs 4B). These data demonstrate that the number of visibly dying neurons in single time point assays is strongly dependent on the rate of microglial efferocytosis, and manipulations that alter the rate of efferocytosis can be misinterpreted as neuroprotective or neurotoxic (Figure 1).

**Fig. 9.**
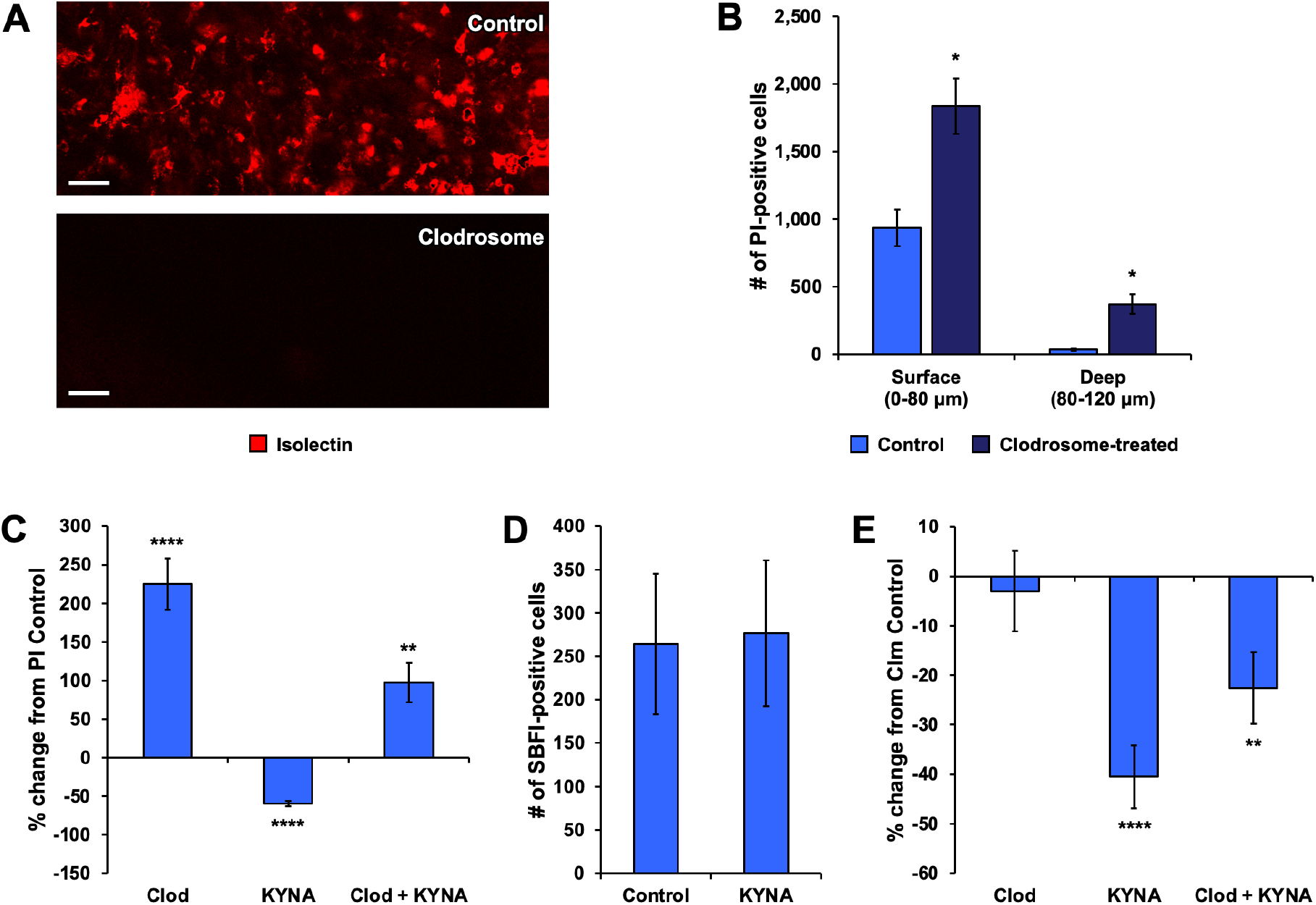
Altering microglial activity affects the number of visible dying neurons. **(A)** Chronic incubation for a minimum of 3 days with liposomal Clodrosome depleted slices of microglia. Scale bars = 50 μm. **(B)** Microglial depletion results in more PI-positive neurons. Clodrosome-treated slices had more PI-positive neurons between 0-80 μm deep (1,836.53 ± 206 vs. 936.17 ± 134, p = 0.0208) and between 80-120 μm deep (373.35 ± 72.30 vs. 34.83 ± 6.37, p = 0.0123). n = 23 slices. **(C)** Kynurenic acid reduces the number of visible dying neurons. Chronic application of 3 mM of KYNA for 10-12 days to block seizures resulted in a reduction in the number of PI-positive neurons compared to control slices (−59.37 ± 3.51 %, p < 0.0001). Chronic application of 0.2 mg/mL of Clodrosome resulted in an increase in the number of PI- positive neurons compared to control slices (224.97 ± 33.24 %, p < 0.0001), as did combined application of KYNA and Clodrosome (97.13 ± 25.27 %, p = 0.0023). n = 53 slices. **(D)** KYNA did not affect the total number of neurons undergoing apoptosis, assayed by SBFI-AM staining. Chronic application of 3 mM of KYNA had no effect on the number of SBFI-positive neurons (n = 23 slices). **(E)** Kynurenic acid reduces the number of visible healthy neurons. Chronic application of 3 mM of KYNA for 10-12 days caused a reduction in the number of Clm-positive neurons compared to control slices (−40.52 ± 6.36 %, p < 0.0001), as did combined application of KYNA and 0.2 mg/mL Clodrosome (−22.57 ± 7.20 %, p = 0.0086). Chronic application of Clodrosome by itself had no effect over the same time period. n = 53 slices.

Based on PI staining, we had concluded in our prior study (38) that seizures were neurotoxic. However, it has recently been reported that neuronal activity (52) and seizures (53) reduce the rate of microglial efferocytosis. To test whether the effects of seizure suppression on the number of PI-positive neurons were due to changes in the rate of microglial efferocytosis, seizures were suppressed in control hippocampal slice cultures and in hippocampal slice cultures in which microglia had been depleted by Clodrosome. DIV 10-12 Clm slices were continuously treated with 3 mM kynurenic acid (KYNA) following the first culture media change on DIV 3, in order to suppress seizure activity (38). Identical Clm slices were treated with 0.2 mg/mL Clodrosome to deplete the slices of microglia, and other slices received both KYNA and Clodrosome. All slices were incubated with PI prior to imaging. Cell counts for a set of slices made from a given mouse pup were normalized to the counts for the untreated control slice(s) from that pup, expressed as percentages, then pooled together with the normalized values from slices from all other pups.

The number of PI-positive neurons was significantly affected by these treatments (Figure 9C). Seizure suppression by KYNA reduced the number of PI-positive neurons compared to control slices (−59.37 ± 3.51 %, p < 0.0001). Depletion of microglia by Clodrosome increased the number of PI+ neurons compared to controls (+224.97 ± 33.24 %, p < 0.0001), as did combined seizure suppression and microglial depletion by application of KYNA and Clodrosome (+97.13 ± 25.27 %, p = 0.0023) (n = 53 slices). These findings indicate that suppressing seizure activity reduces the number of neurons that are PI+ at any given time. This could occur either by reducing the number of neurons entering the apoptotic pathway (i.e. a neuroprotective effect) or by accelerating the efferocytosis of apoptotic neurons via enhanced microglial activity (a false positive neuroprotective effect; Figure 1) (52, 53). The size of the increase in PI+ cells when both seizures and efferocytosis were blocked by KYNA (+97%) was between the reduction in PI+ cells by seizure block alone (−59%) and the increase in PI+ cells by microglial depletion (+225%). Thus microglial depletion reduced the apparent neuroprotective effect of stopping seizures, supporting the idea that stopping seizures increases efferocytosis rather than reduces neuronal apoptosis. However, KYNA reduced PI counts roughly by half in both control and microglia-depleted conditions (a 59% reduction in control conditions, and a reduction from a 225% increase to a 97% increase in Clodrosome). Thus there is a potential microglia-independent effect of KYNA, so that we could not rule out an additional neuroprotective effect of stopping seizures. The difficulty in separating these effects is largely due to the large effect of microglial depletion on the PI counts, i.e. due to microglial phagocytosis being rate-limiting in these experiments. We therefore turned to additional biomarkers to assess these effects.

In slices ranging in age from DIV 2 to DIV 14 there was no difference in the number of neurons which took up SBFI-AM between control slices and age-matched slices in which seizures were blocked by long-term incubation in KYNA starting on DIV 1 (264.27 ± 81.04 vs. 276.33 ± 83.96, n = 23 slices, p = 0.9189; Figure 9D). Because SBFI staining is present throughout apoptosis, this finding suggests that seizures did not alter the balance of neurons entering and exiting the apoptotic pathway. But that interpretation does not seem to be consistent with the reduction in PI+ neurons when seizures were blocked with kynurenate (38) (Figure 9C), if the reduction in PI+ neurons reflects enhanced exit via efferocytosis when seizures are blocked (52, 53). We found however that blocking seizures also *enhanced* entry into the apoptotic pathway, evidenced by an increased rate of extinction of fluorescence of transgenically expressed fluorophores (Figure 9E). Chronic application of 3 mM KYNA for 10-12 days resulted in a *decrease* in the number of Clm-positive neurons by −40.52 ± 6.36 % (p < 0.0001), and KYNA + Clodrosome resulted in a decrease of −22.57 ± 7.20 % (p = 0.0086) (n = 53 slices). The application of Clodrosome without KYNA had no effect over the same time period, indicating that the increased PI+ count in Clodrosome-treated slices is not due to Clodrosome being neurotoxic. This data indicates that prolonged seizure suppression via blockade of ionotropic glutamate receptors can adversely affect healthy neurons, but that a dearth of microglia has no such effect. The difference in survival of healthy FP+ neurons in KYNA was not significantly different than in KYNA + Clodrosome (81).

In addition to reducing the number of Clm-positive neurons, chronic KYNA application also reduced the intensity of Clm fluorescence in surviving neurons (Supplemental Figure 8A, 8B), as would be expected if seizure suppression were adversely affecting healthy neurons. There was also no indication that suppressing seizures with chronic tetrodotoxin (TTX) application for 11-15 days promoted neuronal survival compared to Control conditions when using Clm fluorescence extinction as a metric (Supplemental Figure 8C).

Thus the consistent SBFI-AM staining when seizures are blocked (Figure 9D) reflects both the increased entry into the apoptotic pathway (Figure 9E) and the increased rate of exit (Figure 9C). In contrast to our original interpretation of one-time PI staining data (38), in which we concluded that seizures were neurotoxic, we find that blocking seizures enhances entry into the apoptotic pathway, presumably as a result of reduced neuronal activity (82, 83). Blocking seizures also accelerated exit from the apoptotic pathway by enhancing microglial efferocytosis (52, 53). This is evidenced by the block of the apparent neuroprotective effect of kynurenate (decreased number of PI+ neurons) in the slices depleted of microglia by Clodrosome (Figure 1 & Figure 9C)

These findings indicate that the number of visibly dying neurons is dependent upon the number of microglia available to consume those neurons (Supplemental Figure 6) as well as the phagocytic efficiency of the microglia that consume them (Figure 9). We next tested whether altering the membrane permeability of neurons already committed to apoptosis would alter the rate of biomarker positivity, thereby producing spurious neurotoxic or neuroprotective signals.

### Acute ethanol exposure in the developing hippocampus

Ethanol (EtOH) and other alcohols increase the permeability of biological membranes (84, 85) and so are used to increase access of stains to the cytoplasm of living cells (86, 87) as well as to permeabilize cell membranes after fixation for immunocytochemical (88) and in situ hybridization studies (73). Ethanol consumption in the first 2 trimesters of pregnancy is also clearly linked to human fetal alcohol syndrome (89), which can include microcephaly and developmental disability in addition to dysmorphic facies and behavioral disorders (90). A well-established line of evidence for the neurotoxicity of alcohol is based on one-time assays of biomarkers of apoptosis after alcohol exposure, but the experimental exposure was maximally neurotoxic at later stages of development than occurs in human fetal alcohol syndrome (91). These ethanol toxicity studies utilized the DeOlmos silver stain, a protocol that was developed to sparsely stain degenerating axons, and as such does not include detergent exposure for membrane permeabilization (92). We tested whether ethanol at the clinically relevant concentrations used in these studies could alter the membrane permeability of biomarkers of apoptosis to produce a false positive neurotoxic signal.

DIV 5-7 Clm slices, as well as WT slices infected with AAV9-hSyn-EGFP, were incubated with PI for 60 minutes, then imaged, then incubated with 100 mM EtOH for 24 hours, then incubated with PI again and imaged again. Control slices showed very little change in PI labeling at any slice depth after 24 hours (Figure 10A & C). Conversely, at depths from the sliced surface that were less damaged by slicing (36), 24 hours of ethanol exposure dramatically increased the labeling of neurons for PI (Figure 10B & C). These results suggest that neurons at relatively early stages of apoptosis became more permeable to PI during EtOH exposure. Consistent with this mechanism, cytoplasmic sodium increased dramatically in EtOH-exposed SBFI+ neurons that had had physiological sodium concentrations, but not in EtOH-exposed neurons in late stages of apoptosis whose membranes were already permeable and whose cytoplasmic sodium was consequently already very high (Figure 10D). Acute application of 100 mM EtOH for 60 minutes increased [Na^+^]_i_ by 266.84 ± 65.41 % in neurons with an initial [Na^+^]_i_ between 0-15 mM (n = 68 cells, p < 0.0001), 59.29 ± 15.83 % in neurons with an initial [Na^+^]_i_ between 15-50 mM (n = 73 cells, p = 0.0007), and had no effect on neurons with an initial [Na^+^]_i_ between 50-100 mM (n = 54 cells, p = 0.0621) (n = 10 slices) (Figure 10D).

**Fig. 10.**
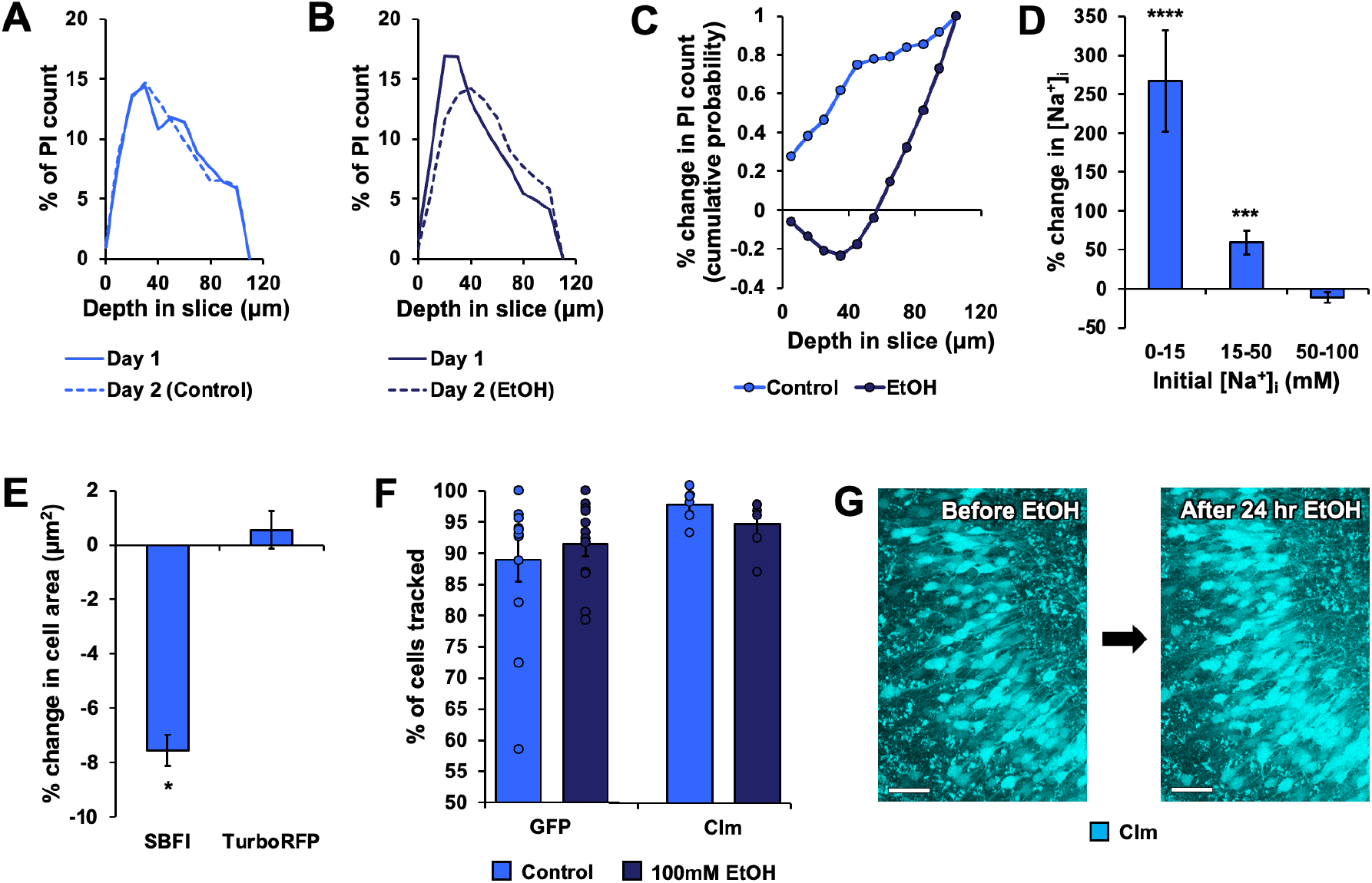
Effects of ethanol on apoptotic and healthy neurons. **(A)** 24 hrs had no effect on the distribution of PI-positive neurons across the entire depth range of Clodrosome-treated slices. n = 8 slices. **(B)** Application of 100 mM ethanol for 24 hrs in Clodrosome-treated slices shifted the distribution of PI-positive neurons towards the deeper regions. n = 9 slices. **(C)** Comparison of data from (A) and (B). 24 hrs of ethanol exposure in Clodrosome-treated slices caused a significant shift in PI-positivity compared to Controls, with PI-positivity decreasing near the surface and increasing in deeper regions. p < 0.0001. n = 8 Control slices and 9 ethanol slices. **(D)** Ethanol application increased [Na^+^]_i_ in apoptotic neurons. Acute application of 100 mM ethanol for 60 minutes increased [Na^+^]_i_ by 266.84 ± 65.41 % in neurons with an initial [Na^+^]_i_ between 0-15 mM (n = 68 cells, p < 0.0001), 59.29 ± 15.83 % in neurons with an initial [Na^+^]_i_ between 15-50 mM (n = 73 cells, p = 0.0007), and had no effect on neurons with an initial [Na^+^]_i_ between 50-100 mM (n = 54 cells, p = 0.0621). n = 10 slices. **(E)** Ethanol application causes shrinkage in apoptotic neurons. Acute application of 100 mM ethanol for 90 minutes caused a 7.56 ± 0.56 % decrease in cell area in SBFI-positive neurons (n = 224 cells from 5 slices, p < 0.0001), but caused no change in cell area in TurboRFP-positive neurons (n = 128 cells from 4 slices, p = 0.4174). **(F)** Ethanol does not reduce the number of visible healthy neurons. Application of 100 mM ethanol for 24 hours caused no reduction in the percentage of GFP- or Clm-positive neurons that could be tracked using TrackMate (GFP: 88.95 ± 3.54 % vs. 91.44 ± 1.95 %, n = 12 slices; Clm: 96.98 ± 1.09 % vs. 93.87 ± 1.70 %, n = 6 slices). **(G)** Example images of Clm-positive neurons before and after 24 hours of 100 mM ethanol exposure, demonstrating little to no quenching or morphological effects. Slice imaged on DIV 7 & 8. Scale bars = 50 μm.

Further supporting the idea of increased membrane permeability in neurons in early stages of apoptosis, but not healthy neurons, were the changes in somatic area in SBFI-positive and TurboRFP-positive neurons (Figure 10E). 90 minutes of 100 mM EtOH exposure caused a 7.56 ± 0.56 % decrease in area in SBFI-positive neurons (n = 224 cells from 5 slices, p < 0.0001), but caused no significant change in area in TurboRFP-positive neurons (n = 128 cells from 4 slices, p = 0.4174).

Finally, the effect of EtOH on the permeability of neurons undergoing apoptosis rather than healthy neurons was supported by the finding that 24 hours of 100 mM EtOH had no effect on healthy neurons expressing the fluorophores GFP or Clm (Figure 10F & G). In EtOH-treated slices, 88.95 ± 3.54 % of neurons in GFP slices and 96.98 ± 1.09 % of neurons in Clm slices survived the treatment and were successfully tracked using TrackMate, compared to 91.44 ± 1.95% and 93.87 ± 1.70 % in Control slices (n = 12 GFP slices and 6 Clm slices). 24 hours of EtOH also had no effect on the number of SBFI-positive neurons in a slice, as the ratio of SBFI+ to TurboRFP+ neurons did not change over that time (Supplemental Figure 9), even as photobleaching due to frequent imaging caused a decrease in TurboRFP fluorescence. This further supports the notion that EtOH application does not result in new neurons entering the apoptotic pathway.

These findings indicate that acute exposure of the perinatal hippocampus to high levels of EtOH increased neuronal labeling with biomarkers of apoptosis by enhancing the permeability of the membrane of neurons undergoing apoptosis rather than damaging healthy neurons. *This data does not indicate that EtOH is not a teratogen*. Rather, these findings indicate that the effects of this experimental protocol primarily reflect a false positive signal: the effects of EtOH on the membrane permeability of neurons undergoing apoptosis, rather than a neurotoxic effect of EtOH on healthy neurons at this developmental stage.

### Prior efferocytosis can create false negatives in cell death analysis

Another source of error in single-time-point analyses of neuronal death can be a failure to account for prior necrosis and efferocytosis when quantifying visible dying neurons. The total amount of neuronal death is not merely the sum of the neurons which stain positive for a cell death biomarker on a given day plus the number that have committed to apoptosis but are not yet positive for the biomarker (Figure 4). Rather, the total amount of neuronal death includes the former categories and in addition, the neurons that have already been ruptured by necrosis, plus the number of apoptotic neurons that had been efferocytosed by microglia prior to the time of analysis.

To illustrate the effect of prior efferocytosis we re-analyzed the data presented in Figure 5A, where slices expressing Clm were incubated once with PI for 60 min on DIV 7 then imaged over several days. There was a clear decrease in PI-positive neurons over a 6-day period (Supplemental Figure 10A). When TrackMate was used to track the initial PI-positive population over time, the number of PI-positive neurons decreased from 546.30 ± 72.88 on DIV 7, to 63.17 ± 18.34 by DIV 10, and finally to 0.33 ± 0.33 by DIV 13 (n = 6 slices) (Supplemental Figure 10B). The 𝛕 of the PI-positive cell loss was calculated to be 1.39 days (Supplemental Figure 10C), which is considerably shorter than the 𝛕 for the loss of SBFI-AM positive neurons (4.69) seen in Figure 7C. This data highlights two key points about the use of PI to assay cell death. First, depending on the timing of the assay relative to injury, quantification of cell death using PI can result in significant underestimation of the total amount of neuronal loss in the slice if PI is only visualized on a single day. In our example, counting PI-positive cells on DIV 10 would only have detected a fraction of the cell death that occurred just 3 days prior. Second, the short 𝛕 for PI cell loss compared to that of SBFI cell loss indicates that neurons become capable of staining positive for PI much later in the apoptotic process than they become capable of staining positive for AM dyes. Thus, utilizing PI without a complementary stain like SBFI that stains neurons earlier in the apoptosis pathway risks further underestimating the total amount of cell death.

## Discussion

### Summary of results

Slicing the brain comprises an acute hypoxic-ischemic and traumatic injury that results in the injury and death of many neurons (36). When brain slices are subsequently cultured, evidence of apoptosis extends into the third week *in vitro* (37, 38). During this time, microglia clear neurons that have entered the apoptotic pathway (Supplemental Figure 6), but there are many more neurons undergoing apoptosis than can be immediately cleared (Figure 4 and Figure 7C). As they await microglial efferocytosis, the permeability of the neurons in the apoptotic pathway increases (Figure 8), resulting in staining for progressively more polar biomarkers of apoptosis (Figure 8H, I). Interventions that alter membrane permeability (Figure 10) or efferocytosis rates (Figure 9) alter the number of visibly dying neurons, and are readily mistaken as being neuroprotective or neurotoxic. Organotypic brain slices thus comprise a useful preparation for the study of neuronal apoptotic death after an acute brain injury.

### Challenges in the assessment of neuroprotection and neurotoxicity

After acute brain injury, the efferocytosis rate is lower than the rate of entry of neurons into the apoptotic pathway (Figure 4). Under these conditions, the assumption that the neuronal death rate is proportional to the number of visibly dying neurons is not valid (Figure 5C). This leads to several potential sources of error in single-time-point assays of neuronal death that we discuss below. The first error is due to necrotic rupture of neurons or efferocytosis of apoptotic neurons prior to the assay for cell death (Supplemental Figure 10) (42). These neurons cannot be counted by the assay and represent a source of false negative error. For example, experimental manipulations that shift apoptotic death to necrotic death could produce a spurious neuroprotective signal by reducing the pool of visibly dying, apoptotic neurons.

The second error is illustrated in Figure 4. One-time assays of neuronal death reveal the size of the pool of neurons that are receptive to the biomarker used. This biomarker is most commonly a stain such as PI, but the biomarker could also be a cellular constituent that is released from dying neurons such as lactate dehydrogenase (24). However, high rates of neuronal death relative to efferocytosis create a queue of neurons that have committed to apoptosis and are awaiting efferocytosis. This queue, in combination with the progressively increasing permeability of the cytoplasmic membrane of the neurons in the queue, can result in shifting, unpredictable inequalities between the population of neurons that are committed to apoptosis and those that are positive for the biomarker. In the case of PI, which stains late in the course of apoptosis (Figure 10I), the associated error can be very large (Figure 5D).

If biomarkers are used that are only transiently positive such as indicators of caspase activity (11), then a related source of error is the loss of biomarker positivity prior to efferocytosis. In this case, the loss of biomarker positivity can be thought of as another way to exit the pool of biomarker-receptive neurons, i.e. another contributor to the rate of exit from this pool in addition to microglial efferocytosis.

A third source of error associated with assays of neuroprotection and neurotoxicity is the rate of microglial efferocytosis relative to the neuronal death rate. If the efferocytosis rate is higher than the rate at which neurons are entering the apoptotic queue, then the pool of neurons receptive for apoptosis biomarkers will shrink by the product of the difference in entry vs exit rates and the length of time over which the rates differ. If the rate of efferocytosis is below the rate at which neurons are entering the biomarker-receptive pool, then the pool will grow as the product of the difference in entry and exit rates and the length of time the rates differ (Figure 4C). Time and experimental manipulations (e.g. Figure 9) can change the rates of entry and exit.

The unanticipated effects of experimental manipulations on the rate of microglial efferocytosis was most clearly demonstrated by microglial depletion, which dramatically reduced microglial efferocytosis rates and increased the size of the biomarker-receptive pool (Figure 9B,C). However, more subtle manipulations of microglial activity such as increasing the microglial efferocytosis rate by inhibition of seizures (53) also had large effects on the size of the pool of biomarker-receptive neurons. Initially we and others have attributed the reduction in the pool of biomarker-receptive neurons when seizures were blocked or absent as a sign that seizures were neurotoxic (37, 38, 49, 50). While the neurotoxicity of sustained seizures in mature animals is considered to be well-established experimentally (93, 94), toxicity is more difficult to ascertain in immature animals (95) and clinically (96–98). We found minimal evidence of such toxicity in this preparation using the emission of transgenically expressed fluorescent proteins as a longitudinal biomarker of neuronal health (Figure 5F; Figure 9E). In fact, blocking seizure activity using the broad spectrum glutamate antagonist kynurenate had the opposite effect on the health of neurons identified by robust fluorescent protein emission (Figure 9E), as has been identified in studies using other broad-spectrum inhibitors of neuronal activity such as the sodium channel antagonist tetrodotoxin (Supplemental Figure 8C) (99).

A fourth error can arise due to unanticipated effects of experimental manipulations on the rate of entry into the pool of neurons receptive to biomarkers of apoptosis. For example, EtOH increases membrane permeability (74, 84, 85, 87, 100) and thereby increased both the cytoplasmic sodium as well as the number of neurons positive for PI, without affecting the number of healthy fluorescent neurons (Figure 10). Because AM dyes permeate early in apoptosis (Figure 6B) and healthy cells did not enter the apoptotic pathway as a consequence of the EtOH exposure (Figure 10E-G), the number of AM dye-positive neurons was not affected by the change in membrane permeability induced by EtOH. Thus, AM dyes are less prone to this error. However, AM dyes may be subject to other sources of error, such as staining neurons or other cells that are not apoptotic, so confirmation with a second method such as fluorescent protein emission is recommended. There is no question that EtOH is a teratogen, and that human fetal alcohol syndrome is characterized by behavioral and cognitive disabilities due to developmental brain injury (89, 90). However, these effects are most strongly associated with exposure in the first two trimesters (89), not the third trimester, which is the stage of human brain development corresponding to the rodent brains used in these experiments near term. We found no evidence of acute EtOH toxicity based on the emission of transgenic fluorescent proteins in healthy neurons in the organotypic hippocampal slice preparation (Figure 10E-G).

### Limitations of this study

This study was carried out in the in vitro organotypic hippocampal brain slice culture preparation in which apoptotic neuronal death after injury was prominent, expression of transgenic fluorophores was facile, and longitudinal multiphoton microscopic evaluation was feasible. The culture media contains a variety of substances such as corticosteroids and insulin that have been empirically found to enhance neuronal survival. We have thoroughly parsed these media components to define their impact on neuronal survival (51), but nevertheless these studies should be replicated both *in vitro* and *in vivo*. For example, hematogenous phagocytic cells were not able to participate in this preparation, although local microglia provide the bulk of phagocytic activity after hippocampal injury (20, 101). As another example, systemic factors that may contribute to the neurotoxicity of prolonged seizures would not be present in the organotypic slice preparation.

The injury that induced the wave of apoptosis in the organotypic slices is a combination of hypoxic-ischemic injury associated with the sacrifice of the rodent as well as trauma associated with brain slicing (36). Other mechanisms of brain injury such as occlusion of arteries or veins, infection, or diffuse traumatic closed head injury may produce different patterns of neuronal death, and these need to be investigated in future studies.

### Implications

The errors associated with assays of neurotoxicity and neuroprotection identified in this study (Figures 1 and 4) can dramatically impact the interpretation and subsequent translation of experimental interventions. For example, based on PI staining we concluded (38) that seizures were killing neurons, when in fact using longitudinal assessment of neuronal health by fluorescent protein emission in this preparation, the opposite was true: blocking seizures killed neurons (Figure 9). The EtOH experiments provide another example of the possibility of misinterpretation of one-time assessments of neurotoxicity (Figure 10) (91). While it may seem clear from clinical data that the overall conclusions of both these experimental studies must be correct, this conclusion bears more thought.

In the case of seizures, the classic studies indicating that prolonged seizures were harmful were performed using convulsants that may have independently killed neurons (94, 102). Human studies linking prolonged seizures with poor outcome are confounded by multiple independently injurious etiologies (96–98), as well as multiple anticonvulsant therapies, which may be neurotoxic (Figure 9). Certainly, prolonged convulsive activity associated with hypopnea, hypoxia, acidosis, and muscle failure is harmful (93). Electrographic seizure activity is used therapeutically (103), so whether electrographic seizures are harmful independently of convulsive activity and the process that incited them is a more complex question that bears continued assessment (104, 105).

In the case of EtOH, there is no question that EtOH is harmful during early stages of development (89, 90), but this fact may bias experimental interpretation (106), as in our study of seizure toxicity (38). An increase in apoptosis biomarkers was evident after neuronal EtOH exposure in perinatal organotypic slice cultures (Figure 10A-C). However, the apparent increase in apoptosis was more likely due to increased membrane permeation of the biomarker, because no direct toxic effect was observed using an assay independent of membrane permeability, i.e. fluorescence emission of transgenic proteins (Figure 10E-G). There is substantial developmental apoptosis in the developing brain whose detection could be magnified by the effects of EtOH on membrane permeability. The experimental paradigm for EtOH toxicity was subsequently extended to anesthetics (107, 108) and anticonvulsants, many of which were administered in solvents such as 10% dimethylsulfoxide (109). The anesthetics and solvents may have enhanced membrane permeability (110, 111) and thus biomarker penetration, as for EtOH. Clinical care has been altered in light of the potential effects of anesthetics and anticonvulsants on developing neurons (112). Thus it will be useful to reassess the neurotoxicity of these agents by longitudinal assessment of neuronal health using transgenic fluorescent protein emission (43).

One of the major challenges of neuroscience is the translation of basic science findings to clinically meaningful therapies (113, 114). The inefficacy of first-generation neuroprotective therapies (115) led to the hypothesis that translational failures arose from deviations from optimal scientific method (116). This led to many important process changes to improve scientific rigor (115, 117–120). The problems identified here, systematic misinterpretation of experimental assays, has not yet been identified as a cause of results that are reproducible but still fail translation (120–122), although recommendations to use multiple outcome measures could reduce the impact of this issue (115). The current findings suggest that assays of neurotoxicity and neuroprotection based on single-time-point assays of neuronal death are easily misinterpreted when neuronal death rates are high, and these misinterpretations could contribute to difficulties with the translation of experimental findings regarding neurotoxicity and neuroprotection. Addition of longitudinal measurements of neuronal health (43) would substantially increase the accuracy of assessments of neuroprotective and neurotoxic effects.

## Materials and Methods

### Culture of organotypic hippocampal slices and experimental conditions

Experiments were performed on C57/BL6 WT and CLM1 (Clomeleon) (from Duke University Medical Center, Durham, NC) mice. Transverse 400 µm hippocampal slices were cut at postnatal day 6-7 (P 6-7) on a McIlwain tissue chopper (Mickle Laboratory Engineering) and cultured using the rocking plate technique (123) or the membrane insert technique (124). Slices were transferred to membrane inserts (PICMORG50; MilliporeSigma) or coverslips, which were placed in glass-bottomed six-well plates (P06-1.5H-N, CellVis) and incubated in 1000 L of NeuroBasal/B27(1x) medium (ThermoFisher Scientific) supplemented with 0.5 mM GlutaMAX and 30 μg/mL gentamicin (both from Invitrogen), in a humidified 37°C atmosphere that contained 5% CO_2_. Culture medium was changed by-weekly. For imaging experiments which required perfusion, slices were transferred to a submerged chamber and continuously superfused in oxygenated (95% O_2_, 5% CO_2_) ACSF containing: 126 mM NaCl, 3.5 mM KCl, 2 mM CaCl_2_, 1.3 mM MgCl_2_, 25 mM NaHCO_3_, 1.2 mM NaHPO_4_, and 11 mM glucose (pH 7.4). ACSF was perfused at 32°C and recirculated for the duration of the imaging session, unless otherwise noted. Organotypic hippocampal slices were used at DIV 1-40.

Pharmacological agents included: KYNA at 3 mM to suppress seizure activity, ouabain at 10 μM to block Na/K ATPases, 10Panx at 50 μM to block pannexin-1 hemichannels, liposomal clodronate (Clodrosome) at 0.2 mg/mL to eliminate microglia, and EtOH at 100 mM. All pharmacological agents were from MilliporeSigma, except for Clodrosome which was from Encapsula NanoSciences.

### Imaging

Two-photon imaging was performed using a custom-built scanning microscope. Two-photon images were acquired using custom-designed software (LabVIEW), a scan head from Radiance 2000 MP (Bio-Rad) equipped with a 40x, 0.8 NA water-immersion objective (Olympus), and a mode-locked Ti:Sapphire laser (MaiTai; Spectra-Physics, Fremont, CA). Excitation wavelengths varied by experiment. Emission was detected through three filters: 470/50 nm, 545/30 nm, and 620/100 nm. These were optimized for CFP, YFP, and RFP emission but were also used to visualize other fluorophores with blue/green, yellow, and red emission respectively. Three photomultiplier tubes (Hamamatsu Photonics) were used to acquire signals in all three emission ranges simultaneously when necessary. Three-dimensional stacks (3D) of raster scans in the XY plane (0.92 μm/pixel XY in most cases, 0.61 μm/pixel XY when quantifying intracellular ion concentrations) were imaged at a z-axis interval of 2 μm. Images were reconstructed offline either by ImageJ (RRID: SCR-003070) or MATLAB (MathWorks).

### Fluorophores

SBFI, AM, cell permeant (SBFI; ThermoFisher) was diluted in Pluronic F-127 (20% solution in DMSO) (ThermoFisher) and incubated in slices overnight at a concentration of 27 μM (final DMSO concentration = 0.3%). During imaging SBFI was excited at 725 and then at 800 nm, and emission was detected through the 445-495 nm filter. Fura Red, AM, cell permeant (FuraRed; ThermoFisher) was diluted in Pluronic F-127 (20% solution in DMSO) (ThermoFisher) and incubated in slices overnight at a concentration of 28 μM (final DMSO concentration = 0.3%). During imaging FuraRed was excited at 800 and then at 875 nm, and emission was detected through the 570-670 nm filter.

SBFI and FuraRed were calibrated to allow for the ratiometric quantification of [Na^+^]_i_ and [Ca^2+^]_i_, respectively. For SBFI, regular ACSF and sodium-free ACSF (substituting mannitol for NaCl) were prepared and combined in various ratios, resulting in solutions with sodium concentrations ranging from 0 to 150 mM. In the presence of gramicidin (a perforating agent which equalizes the intracellular and extracellular sodium concentrations), slices loaded with SBFI were imaged while these solutions were perfused in succession. SBFI was excited at 725 and then at 800 nm, and the ratio of the emission intensity resulting from both excitations was calculated for many cells, at each extracellular sodium concentration. The 725/800 ratio was graphed versus the sodium concentration, and a line of best fit was determined. The equation describing this line was then used to derive [Na^+^]_i_ for each cell during future SBFI imaging experiments, given only the emission intensity for that cell when excited at 725 and 800 nm. This calibration protocol was repeated periodically to ensure that any changes to the equipment (regular maintenance, laser realignments, etc.) did not skew the [Na^+^]_i_ values over time. For FuraRed, calibration utilized a Calcium Calibration Buffer Kit (ThermoFisher) containing buffer solutions with free calcium concentrations ranging from 0 to 39 μM with a mix of K_2_-egtazic acid (EGTA), CaEGTA, KCl, and 3-(*N*-morpholino)propanesulfonic acid (MOPS). These solutions were combined at the ratios specified in the kit, and perfused over FuraRed-loaded slices in the presence of ionomycin (to equilibrate extracellular and intracellular calcium). FuraRed was excited at 800 and then at 875 nm, and the 800/875 ratio of emission intensities was graphed versus the free calcium concentration. The line of best fit and resulting equation were then derived and utilized for future experiments, as with SBFI.

Propidium iodide (1.0 mg/mL solution in water) (ThermoFisher) was incubated in slices for 1 hr at a concentration of 6 μM. During imaging PI was typically excited at 800 nm, but where indicated was excited off-peak so as to excite another fluorophore simultaneously for coregistration. PI emission was detected through the 570-670 nm filter. FAM-FLICA from the FAM-FLICA Caspase-3/7 Assay Kit (Green FLICA; ImmunoChemistry) was diluted in 50 μL of DMSO and 5 μL of this stock was added to 1 mL of media to incubate in slices for 30-60 min (final DMSO concentration = 0.5%). During imaging Green FLICA was excited at 850 nm and emission was detected through the 445-495 nm filter. SR-FLICA from the SR-FLICA Poly Caspase Assay Kit (Red FLICA; ImmunoChemistry) was diluted in 50 μL of DMSO and 5 μL of this stock was added to 1 mL of media to incubate in slices for 30-60 min (final DMSO concentration = 0.5%). During imaging Red FLICA was excited at 800 nm and emission was detected through the 570-670 nm filter. Annexin V, Alexa Fluor 594 conjugate (Annexin V; ThermoFisher) was incubated in slices for a minimum of 2.5 hrs at a ratio of 50 μL per 1 mL of media. During imaging Annexin V was excited at 800 nm and emission was detected through the 570-670 nm filter. PO-PRO-1 iodide (435/355) (1 mM Solution in DMSO) (PO-PRO; ThermoFisher) was incubated in slices for a minimum of 3 hrs at a concentration of 5 μM (final DMSO concentration = 0.5%). During imaging PO-PRO was excited at 775 nm and emission was detected through the 445-495 nm filter. NucBlue Live ReadyProbes Reagent (Hoechst 33342) (NucBlue; ThermoFisher) was incubated in slices overnight at a ratio of 4 drops per 1 mL of media. During imaging NucBlue was excited at 775 nm and emission was detected through the 445-495 nm filter. Isolectin GS-IB4 from Griffonia simplicifolia, Alexa Fluor 594 conjugate (Isolectin; ThermoFisher) was incubated in slices for a minimum of 2.5 hrs at a concentration of 10 μg per 1 mL of media. During imaging Isolectin was excited at 800 nm and emission was detected through the 570-670 nm filter.

The AAV9-hSyn-TurboRFP viral vector (TurboRFP; AddGene) was incubated in slices for a minimum of 24 hrs at a ratio of 5 μL per mL of media. During imaging TurboRFP was typically excited at 750 nm, but was sometimes excited off-peak so as to excite another fluorophore simultaneously for coregistration. TurboRFP emission was detected through the 570-670 nm filter. The AAV9-hSyn-eGFP viral vector (GFP; AddGene) was incubated in slices for a minimum of 24 hrs at a ratio of 5 μL per mL of media. During imaging GFP was excited at 800 nm and emission was detected through the 445-495 nm filter. The AAV9.hSyn.HI.eGFP-Cre.WPRE.SV40 viral vector (GFPcre; AddGene) was injected into P4 mouse pups using intracerebroventricular (ICV) injection (125). During imaging GFPcre was excited at 900 nm and emission was detected using the 530-560 nm filter. The AAV5-gfa2-GFP viral vector (GFAP-GFP; University of Pennsylvania), which utilizes a GFAP-derived promoter, was incubated in slices for a minimum of 24 hrs at a ratio of 0.5 μL per mL of media. During imaging GFAP-GFP was excited at 850 nm and emission was detected through the 445-495 nm filter.

During imaging of slices from Clomeleon pups an excitation wavelength of 860 nm was used, and emission was detected through both the 445-495 nm (CFP) and the 530-560 nm (YFP) filters simultaneously.

AAV-syn-jRCaMP1a and AAV-syn-cre-nls-GFP (both from AddGene) were injected into P1 mouse pups using ICV injection. Imaging of these fluorophores utilized a different, single-photon setup called an “incuscope” (40, 58). During imaging the GFP was excited at 459 nm and emission was detected using a 500-550 nm filter.

### Serial imaging experiments

Experiments to assess the degree of overlap of TurboRFP expression and FLICA staining were performed using the submerged perfusion chamber. Rocking plate slices expressing TurboRFP and loaded with FLICA were placed in the chamber with perfusion operating as normal, and imaged to assess the emission intensity of the two fluorophores and overlap between the two. In a subset of these experiments, multiple fields of view (FOVs) were imaged in rapid succession. In instances where serial observations were necessary, after the baseline images of a given FOV were obtained (at “Hour 0”) the perfusion was shut off in order to deprive the neurons of heat as well as fresh oxygen and nutrients. The same FOV (“Slice region 1”) was then imaged once every hour for three more hours. The FOV was then moved to a different region of the hippocampal pyramidal layer (“Slice region 2”) and this new FOV was imaged over three more hours.

Experiments to assess the survival of SBFI- or PI-positive neurons, often alongside neurons expressing a fluorescent protein such as TurboRFP or Clm, were performed in two ways. When a small number of time points were required over a period of 24 hours, rocking plate slices were imaged, returned to the incubator, then imaged again the following day. When a higher degree of temporal resolution over a longer period of time was required, we employed a TC-MIS miniature incubator connected to a TC-1-100-I temperature controller (both from Bioscience Tools). Slices on membrane inserts were placed inside the mini-incubator, the humidified atmosphere of which was set to match the conditions that the slices experience during normal incubation (37°C and 5% CO_2_). Under these conditions, slices could survive for extended periods of time until the fourth day, when the slice began to suffer the effects of not having their culture medium changed. Slices were imaged using the same microscope, laser, and objective (mounted to an inverter arm positioned beneath the mini-incubator) as in all other experiments. For serial observations of PI + FP, Clm slices or slices expressing TurboRFP were incubated with PI for one hour prior to imaging, then washed, then placed inside the mini-incubator. For serial observations of SBFI + FP, slices were incubated with SBFI overnight prior to the start of the experiment, and SBFI was present in the media during the experiment (to allow for the appearance of newly SBFI-positive cells over time). Images were acquired as frequently as every hour, though typically every two hours proved sufficient. This mini-incubator setup was employed for experiments > 1 day in total duration (to minimize the chances of bacterial contamination) and for experiments where > 3 hrs of continuous imaging was necessary (to better maintain the health of the slices).

Experiments using isolectin to visualize microglial engulfment of SBFI-positive neurons also employed the mini-incubator setup. Imaging with high temporal resolution was necessary to visualize the rapid changes in positioning and morphology common with active microglia. Slices expressing GFP from the AAV-syn-cre-nls-GFP virus were imaged using a microscope and LED light source mounted inside an incubator, and could be continuously perfused and serially imaged. This “incuscope” setup was described previously (40, 58).

All figures which present images taken at multiple time points from the same experiment use identical acquisition and post-processing parameters.

### In vivo experiments

Craniotomies were performed on postnatal day 1 to 3 (P 1-3) Clm mice (corresponding to the VLBW gestational ages 23-32 weeks) following previously described procedure (126). Mouse pups were anesthetized with isoflurane (2%) while the skull overlaying the region of interest was removed using a scalpel blade. A thin layer of 1.5% low-melting point agarose (Millipore-Sigma) dissolved in cortex buffer (25 mM NaCl, 5 mM KCl, 10 mM glucose, 10 mM HEPES, 2 mM CaCl_2_, and 2 mM MgSO_4_, pH 7.4) (127) was applied to the exposed dura. The cranial window was sealed with 3 mm diameter cover glass (Warner Instruments) which was secured to the skull using dental cement. A custom-made titanium headbar was attached to the area adjacent to the cranial window.

During imaging, pups were kept under anesthesia with isoflurane (2%) and their body temperature was maintained using a heating pad (Kent Scientific). Focal cerebral ischemia was induced by photothrombosis of cortical blood vessels using Rose Bengal (Millipore-Sigma) (62–64). Prior to imaging, mice received intraperitoneal (IP) injection of 50 mg/kg Rose Bengal and ischemia was induced by focal illumination of the cortex (128). Cerebral reperfusion or occlusion was established using either the intravascular Rose Bengal itself or IP injection of dextran-conjugated (70 kDa) Texas-Red fluorescent dye (129).

### In vitro electrophysiology

DIV 6-14 Clm organotypic slice cultures were loaded overnight with FuraRed, then transferred to a recording chamber and perfused with ACSF (2.5 ml/min) containing the following (in mM): 124 NaCl, 1.25 NaH_2_PO_4_, 2.5 KCl, 26 NaHCO_3_, 2 CaCl_2_, 2 MgSO_4_, and 20 D-glucose, bubbled with 95% O_2_ and 5% CO_2_ at 34°C. Whole-cell patch-clamp recordings were performed on Clm-positive or FuraRed-positive CA1 pyramidal cells, where cells were visualized by an upright microscope (Eclipse FN1, Nikon) equipped with a 40X, 0.8 NA water-immersion objective (Nikon). Electrodes were pulled from borosilicate glass capillaries (Sutter Instruments) using a micropipette puller (model P97, Sutter Instruments) with resistance of 5-7 MΩ when filled with internal solution containing the following (in mM): 124 K-MeSO_4_, 5 KCl, 10 KOH, 4 NaCl, 10 HEPES, 28.5 sucrose, 4 Na_2_ATP, 0.4 Na_3_GTP, and 1.4 mM 6-methoxy-*N*-ethylquinolinium iodide (MEQ), with osmolarity of 295 mOsm and pH 7.25-7.35. Resting membrane potential was measured and compared between Clm and FuraRed-positive neurons after whole-cell configuration was reached. Only neurons with series resistance < 20 MΩ (assessed in voltage-clamp mode by −5 or −10 mV voltage step) were included in the analysis. Signal acquisition was performed using a Multiclamp amplifier (Multiclamp 700B, Molecular Devices) with Clampex 10 software (Molecular Devices). Signals were sampled at 10 kHz and filtered at 2 kHz. Data were stored on a PC for offline analysis after digitization using an analog-to-digital converter (Digidata 1440A, Molecular Devices).

### Image analysis

Images were analyzed predominantly in ImageJ. Fluorescence intensity was obtained by manually drawing a region of interest (ROI) around the cell or nucleus in question. In cases where the cell or nucleus was present in multiple z-axis frames, the frame in which the intensity was greatest was used. Two-dimensional area was similarly obtained by drawing a ROI around the cell or nucleus in the z-frame with the greatest intensity.

Cell counting and tracking was performed using the TrackMate plugin in ImageJ (130). Prior to TrackMate analysis, images in a time series were manually cropped, rotated, and aligned as necessary to control for changes in slice size and position over time, so as to aid in cell tracking. For both spot detection (cells) and track detection (tracking cells over multiple time points) all default TrackMate settings were used with one exception: estimated blob diameter (the approximate diameter of the object being detected) was increased from 10 to 20 μm when counting FP+ neurons to prevent neuronal processes from being incorrectly counted as soma. By default TrackMate uses a Laplacian of Gaussian (LoG) filter to find Gaussian-like particles in the presence of noise. It applies a LoG filter, looks for local maxima, and each detected spot is assigned a Quality value which is larger for bright spots and spots whose diameter is close to the specified diameter. Thus spots were filtered to remove autofluorescence and background fluorescence using a filter for Quality values above a certain threshold, with the threshold being set automatically by the software in most cases. In instances in which the same threshold needed to be preserved across multiple time points, the auto-thresholded Quality value for one time point was manually assigned to all other time points; this exemplar time point was either the first time point in a time series, or the time point where the fluorophore of interest was brightest, depending on the experimental design. Overlap between multiple fluorophores was quantified in MATLAB using the spot detection data obtained from TrackMate.

Determinations of slice health based on autofluorescence were made using CellProfiler. The diameter range for cell detection was set to 10-50 pixels, and for each image the average diameter, area, and perimeter of all detected cells was exported and then grouped by slice.

𝛕 values for the progressive loss of cells in a population were calculated using Microsoft Excel. The percentage of the initial cell population remaining at each time point was plotted on a graph, and an approximate 𝛕 was determined. This 𝛕 was then used to generate an approximate monoexponential decay curve which conformed to the experimental data. The difference between the experimental and approximate curve-derived values was taken for each time point, and the differences for all time points were summed. The Solver plugin was then used to determine the value for 𝛕 that resulted in the smallest sum of differences between the real data and the curve. This 𝛕 was therefore the value that yielded an exponential decay curve that best fit the loss of the initial cell population.

### Statistics

Imaging data was analyzed with ImageJ or MATLAB. Statistical analysis of the data was done in GraphPad Prism (GraphPad Prism 9). Statistics were assessed with two-tailed Student’s t-tests (unpaired or paired, depending on experimental design) when comparing two groups and one-way ANOVAs with repeated measures when comparing more than two groups, unless otherwise noted. The number of data points (n) and the statistical significance (p-value) are stated in figure legends. All error bars in figures are presented as mean ± standard error.

### Study approval

All experiments were performed in accordance with protocols approved by the Center for comparative medicine (CCM) at Massachusetts General Hospital (MGH) and in accordance with the National Institute of Health Guide for the Care and Use of Laboratory Animals.

## Data Availability

Data can be made available upon request.

## Author contributions

Conceptualization: TB, KS

Methodology: TB, KL, NR, FB, VD, KS

Investigation: TB, KL, NR, FB, VD

Visualization: TB, KS

Supervision: KS

Writing—original draft: TB, KS

Writing—review & editing: TB, EB, KS

