## Supplemental Figures for "A dynamic balance between neuronal death and clearance after acute brain injury"

| Plate # | Slice # | Day 1 PI | Day 2 PI | Day 3 PI | Day 4 PI |
|---|---|---|---|---|---|
| **1** | **1** | 180 | 151 | 156 | 131 |
| **1** | **2** | 17 | 25 | 30 | 18 |
| **1** | **3** | 4 | 6 | 3 | 1 |
| **1** | **4** | 26 | 120 | 17 | 82 |
| **1** | **5** | 19 | 34 | 8 | 28 |
| **1** | **6** | 23 | 186 | | |
| **2** | **1** | 4 | 5 | 25 | 27 |
| **2** | **2** | 4 | 2 | 1 | 1 |
| **2** | **3** | 6 | 3 | 3 | 7 |
| **2** | **4** | 3 | 16 | 12 | 13 |
| **2** | **5** | 3 | 1 | 1 | 1 |
| **2** | **6** | 0 | 0 | 35 | 31 |
| **3** | **1** | 6 | 9 | 13 | 10 |
| **3** | **2** | 12 | 0 | 3 | 6 |
| **3** | **3** | 2 | 3 | 2 | 1 |
| **3** | **4** | 27 | 1 | 1 | |
| **3** | **5** | 2 | 6 | 4 | 22 |

**Table S1. Propidium iodide counts from 4-day serial imaging experiment.**
Raw PI counts from the 4-day serial imaging experiment summarized in Figures 5C and 5D. 17 slices from 3 rocker plates were imaged 4 times over 72 hours and the ImageJ plugin TrackMate was used to count PI-positive cells.

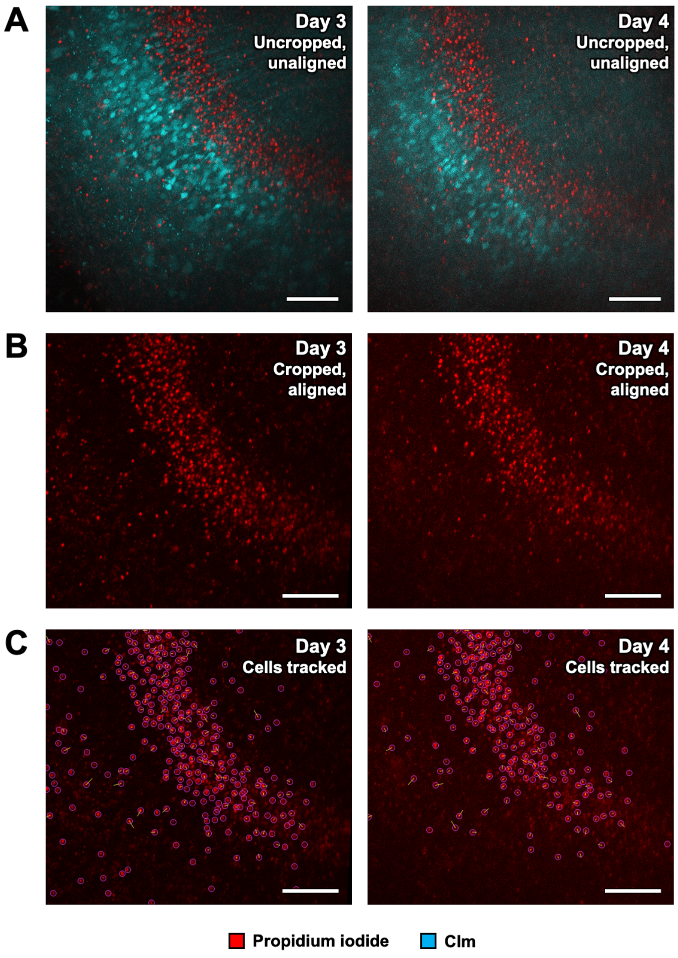


**Fig. S1. Using ImageJ and TrackMate to count and track PI-positive neurons.**
**(A)** Example of Clm + PI merged images. Clm and PI channels were subtracted from each other in TrackMate to minimize background fluorescence and autofluorescence, then the resulting images were merged. Experiment Day 3 = DIV 9. Scale bars = 100 µm. **(B)** Example of aligned PI images. Images from different days were rotated and shifted along the x- and y-axes to facilitate alignment using Clm-positive neurons as landmarks. Scale bars = 100 µm. **(C)** Example of cell counting as tracking using TrackMate. Using the rotated and aligned PI images, the TrackMate plugin was used to count PI-positive cells and track them if they were present across multiple experiment days. Pink circles indicate “spots” (cells) detected, and yellow lines (tracks) indicate movement of cells over multiple days. Scale bars = 100 µm.

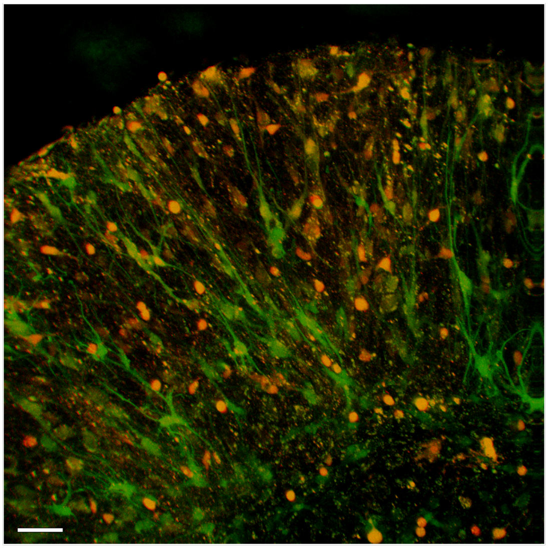

GFAP-GFP SBFI

**Fig. S2. AM dye-positive cells are neurons.**
Example image showing the lack of overlap between SBFI and GFAP-GFP, which is expressed only in astrocytes. Slice imaged on DIV 12. Scale bar = 50 µm.

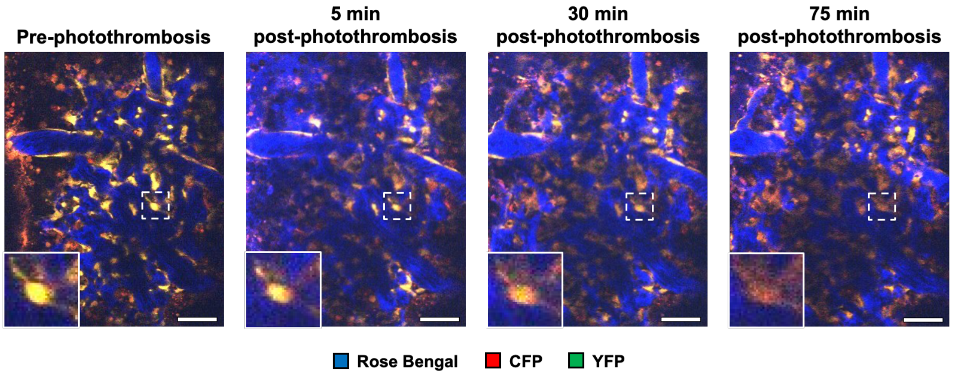


**Fig. S3. Fluorescent protein quenching as a cell death biomarker *in vivo*.**

Example images showing fluorescent proteins quenching as a result of photothrombosis *in vivo*. Anesthetized Clomeleon mice that were subjected to photothrombosis using Rose Bengal experienced widespread dimming of both CFP and YFP over 75 min. Inset: example neuron. Mouse imaged at age P3. Scale bars = 50 µm.

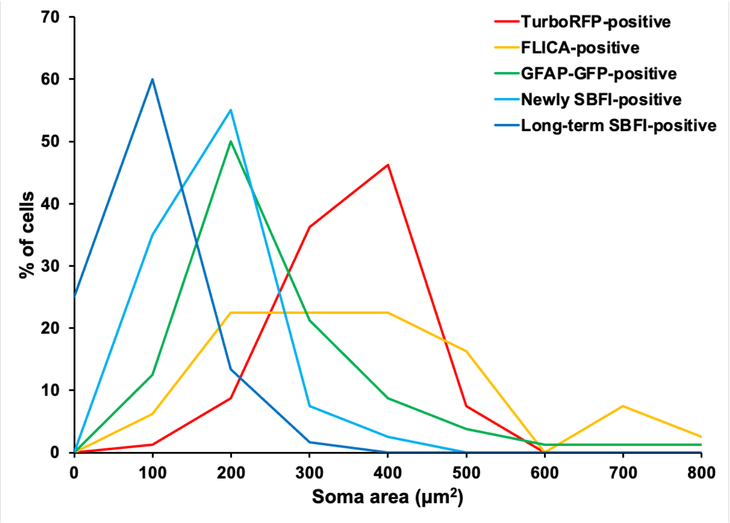


**Fig. S4. Distribution of soma areas for fluorophore-positive cells.**
Histogram showing the distribution of somatic area for cells positive for various fluorophores: TurboRFP-positive cells (n = 80), FLICA-positive cells (n = 80), GFAP-GFP-positive cells (n = 80), cells SBFI-positive for less than 24 hrs (n = 80), and cells SBFI-positive for more than 24 hrs (n = 60).

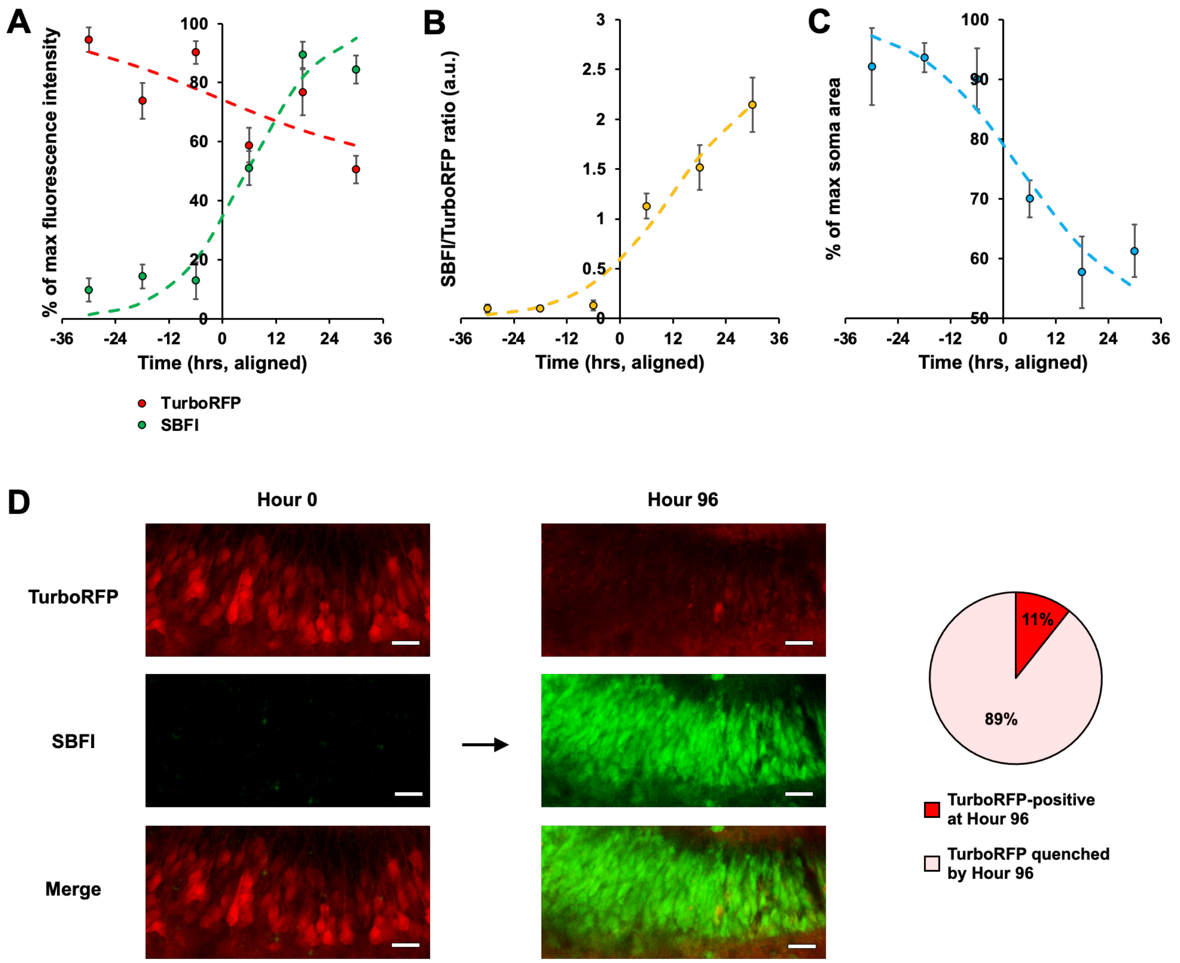


**Fig. S5. Fluorescent protein quenching vs. AM dye uptake.**

**(A)** AM dye uptake and fluorescent protein quenching occur simultaneously. Depriving slices of fresh media for more than a few days resulted in widespread AM dye uptake and fluorescent protein quenching. 15 neurons from 5 slices were visualized in ImageJ, and SBFI and TurboRFP fluorescence intensity were both quantified. The ratio of SBFI intensity to TurboRFP intensity for a given neuron was calculated, the time interval at which this ratio experienced its largest increase was taken to be time = 0 for the observable initiation of cell death, and all other time points for that neuron were aligned accordingly. The largest drop in TurboRFP fluorescence (90.28 ± 3.84 % of maximum to 58.78 ± 5.85 %) and the largest increase in SBFI fluorescence (12.95 ± 6.22 % of maximum to 51.08 ± 5.83 %) were found to occur during the same time interval. **(B)** The SBFI/TurboRFP intensity ratio versus time, as described in (A). The time interval over which the ratio experienced its largest increase (0.13 ± 0.05 to 1.13 ± 0.13) was used to align the time points time = 0 for each neuron individually. **(C)** Soma shrinkage also occurs alongside AM dye uptake and fluorescent protein quenching. The largest drop in 2-D soma area (90.07 ± 5.08 % of maximum to 70.03 ± 3.06 %) occurred during the same time interval as did maximum SBFI uptake and maximum TurboRFP quenching, indicating that all three processes occur concurrently as the cell death process begins. **(D)** Visualization of widespread SBFI uptake and TurboRFP quenching. A slice from the experiments quantified in

(A) through (C), demonstrating plentiful TurboRFP expression and sparse SBFI uptake at Hour 0, contrasted with near-total (89%) quenching of the same TurboRFP-positive neurons Hour 96 and SBFI uptake having become extremely common. ImageJ and TrackMate were used to visualize and track neurons over time. Scale bars = 50 μm.

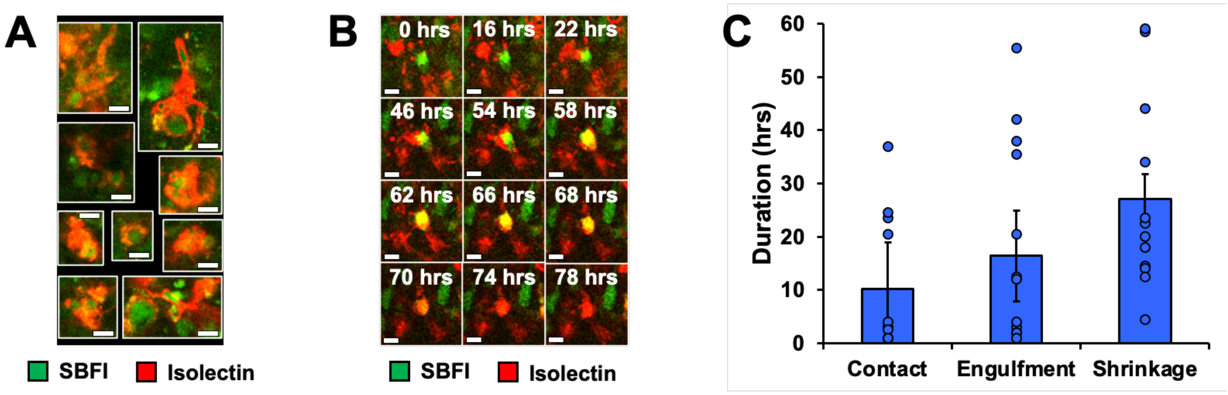


**Fig. S6. Microglial engulfment is the final stage in delayed neuronal death.**

**(A)** Example images of active microglia stained with isolectin surrounding and engulfing SBFI-positive neurons. Scale bars = 10 µm. **(B)** Sequential images showing an SBFI-positive neuron coming into contact with, and ultimately being destroyed by, microglia over 78 hours. Scale bars = 10 µm. **(C)** The duration of microglial engulfment is highly variable. The time during which microglia make contact with SBFI-positive neurons but before they engulf them ("Contact"), the time during which the neurons are engulfed by the microglia but are not yet shrinking ("Engulfment"), and the time during which the neurons are undergoing terminal cell shrinkage while being engulfed ("Shrinkage") all varied widely (n = 18 cells from 2 slices).

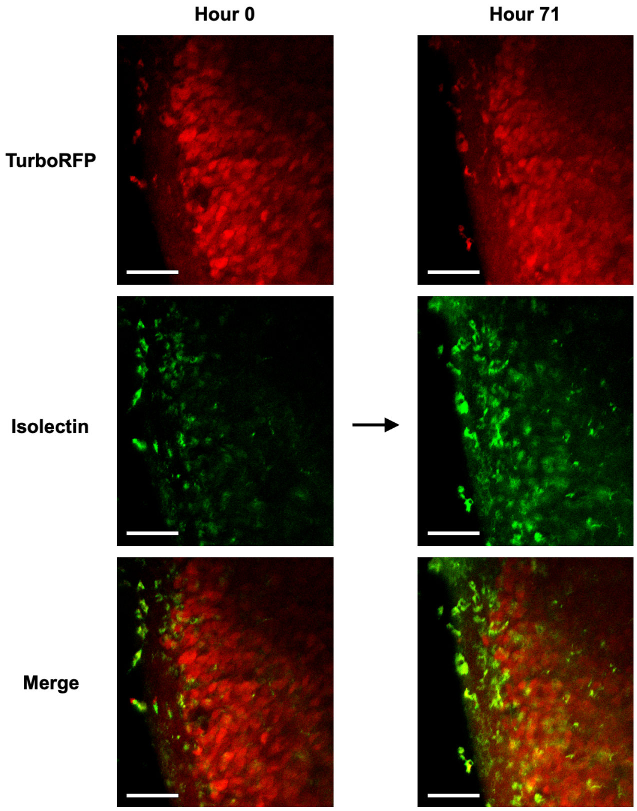


**Fig. S7. Fluorescent protein-positive neurons are not targeted by microglia.**
In a serially-imaged membrane slice with neurons expressing TurboRFP and active microglia stained with isolectin, no instances of microglial engulfment of neurons could be seen over the course of 3 days, indicating that efferocytosis is limited to neurons which have already entered the apoptosis pathway. Experiment Day 0 = DIV 12. Scale bars = 100 μm.

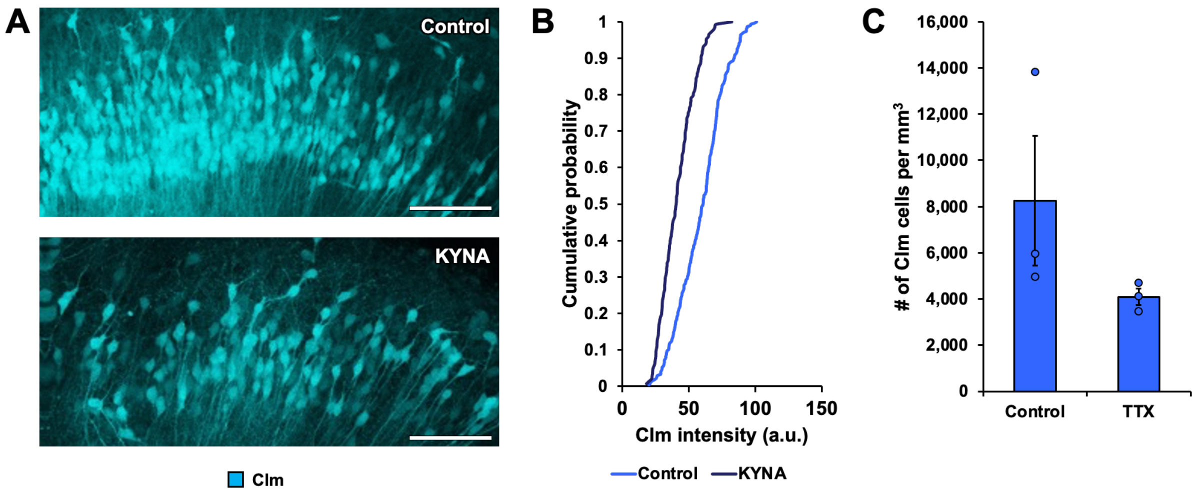


**Fig. S8. Clm fluorescence in Control and seizure-blocked slices.**

(A) Example images of a Control slice and a slice that was treated with KYNA for 10-12 days, used for the analysis in Figures 9C-E. KYNA-treated slices have fewer visible Clm-positive neurons. Slices imaged on DIV 10. Scale bars = 100 µm. **(B)** Quantification of the fluorescence emission intensity of Clm-positive neurons in Control vs. KYNA-treated slices. Neurons in KYNA-treated slices have a lower overall emission intensity. Clm intensity is presented as the 3$^{rd}$ quartile, or 75$^{th}$ percentile, instead of the mean in order to minimize the influence of dark background pixels included in the ROIs. **(C)** Quantification of visible Clm-positive neurons in Control vs. TTX-treated slices. There was no indication that treatment with TTX for 11-15 days increased neuronal survival over Control slices.

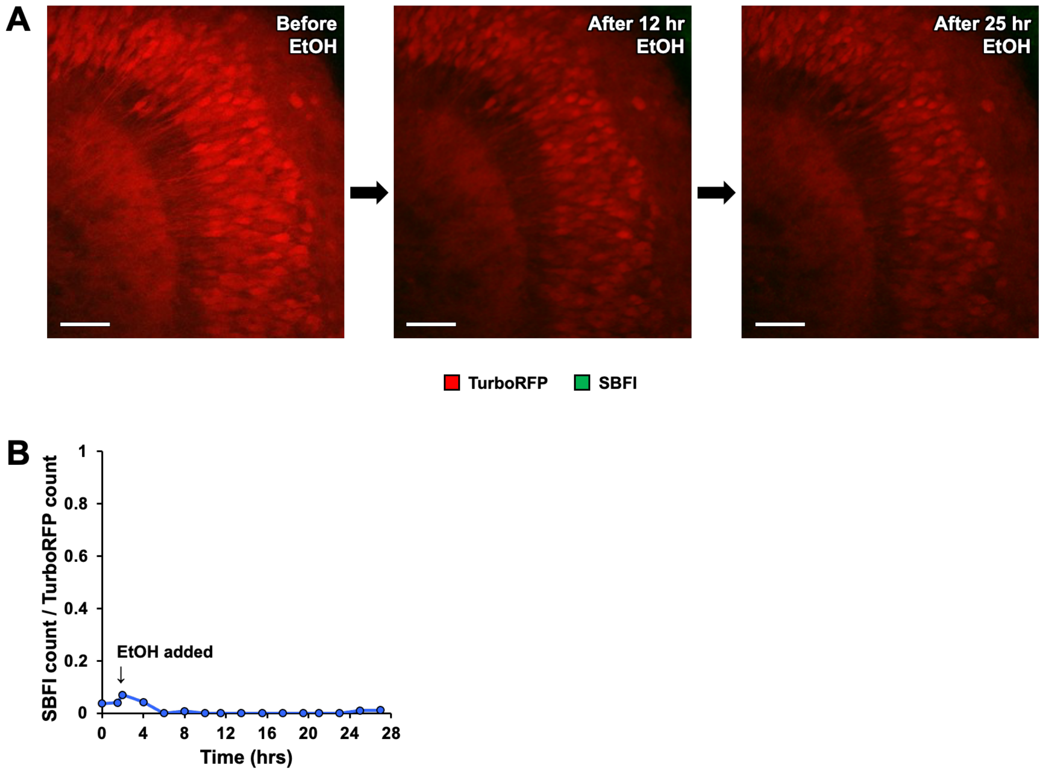


**Fig. S9. Ethanol application does not increase the number of AM dye-positive cells.**

**(A)** EtOH does not cause additional AM dye uptake. In a DIV 12 slice expressing TurboRFP and incubated with SBFI, 100 mM EtOH was added after obtaining a 2 hr imaging baseline, and the slice was imaged every 2 hrs hence. No new SBFI-positive neurons were visually apparent, even after >24 hrs. Due to the frequent imaging some TurboRFP photobleaching was seen, but this was not TurboRFP quenching from the EtOH application since the fluorescent protein emission dimmed somewhat then remained constant, instead of continuing to dim until it was no longer visible. Scale bars = 100 μm. **(B)** Quantification of the lack of new AM dye uptake. For >24 hrs after 100 mM EtOH application, there was no increase in the ratio of SBFI+ neurons to TurboRFP+ neurons.

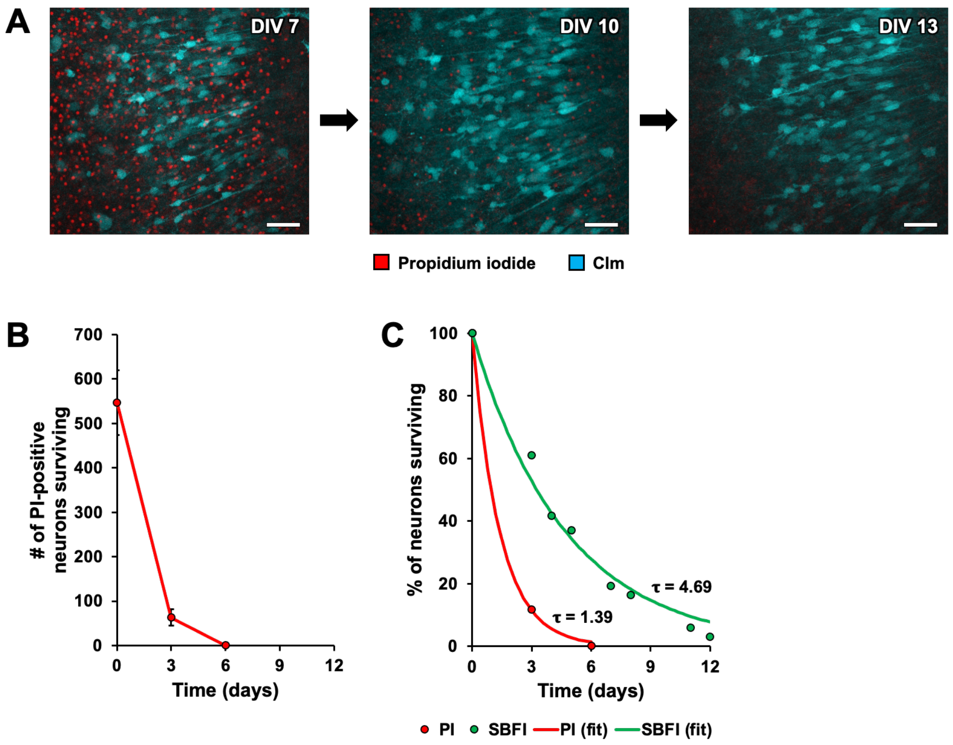


**Fig. S10. Prior phagocytosis can create false negatives in cell death analysis.**

**(A)** Apoptotic neurons disappear over time. Slices expressing Clm were given a single dose of PI on DIV 7, then imaged over several days. There was a marked decrease in PI-positive neurons over a 6-day period. Scale bars = 50 µm. **(B)** Quantification of the apoptotic cell loss. TrackMate was used to track individual cells over time. From 546.30 ± 72.88 PI-positive cells on DIV 7, the count decreased to 63.17 ± 18.34 cells by DIV 10, and 0.33 ± 0.33 cells by DIV 13 (n = 6 slices). **(C)** SBFI-positive cells survive for much longer than PI-positive cells. The time constant (**τ**) of the PI-positive cell loss (1.39) is much shorter than the **τ** for the loss of SBFI-positive cells (4.69), as previously described in Figure 7C.
